# Genetic and functional diversity help explain pathogenic, weakly pathogenic, and commensal lifestyles in the genus *Xanthomonas*

**DOI:** 10.1101/2023.05.31.543148

**Authors:** Michelle M. Pena, Rishi Bhandari, Robert M. Bowers, Kylie Weis, Eric Newberry, Naama Wagner, Tal Pupko, Jeffrey B. Jones, Tanja Woyke, Boris A. Vinatzer, Marie-Agnès Jacques, Neha Potnis

## Abstract

The genus *Xanthomonas* has been primarily studied for pathogenic interactions with plants. However, besides host and tissue specific pathogenic strains, this genus also comprises nonpathogenic strains isolated from a broad range of hosts, sometimes in association with pathogenic strains, and other environments, including rainwater. Based on their incapacity or limited capacity to cause symptoms on the host of isolation, nonpathogenic xanthomonads can be further characterized as commensal and weakly pathogenic. This study aimed to understand the diversity and evolution of nonpathogenic xanthomonads compared to their pathogenic counterparts based on their co-occurrence and phylogenetic relationship and to identify genomic traits that form the basis of a life-history framework that groups xanthomonads by ecological strategies. We sequenced genomes of 83 strains spanning the genus phylogeny and identified eight novel species, indicating unexplored diversity. While some nonpathogenic species have experienced a recent loss of a type III secretion system, specifically, the *hrp2* cluster, we observed an apparent lack of association of the *hrp2* cluster with lifestyles of diverse species. We gathered evidence for gene flow among co-occurring pathogenic and nonpathogenic strains, suggesting the potential of nonpathogenic strains to act as a reservoir of adaptive traits for pathogenic strains and vice versa. We further identified traits enriched in nonpathogens that suggest a strategy of stress tolerance, rather than avoidance, during their association with a broad range of host plants.

## Introduction

The genus *Xanthomonas*, traditionally considered to group plant pathogenic bacteria, encompasses bacterial strains that although they maintain close association with plants, do not cause apparent disease symptoms in their host of isolation (Bansal et al., 2021; Essakhi et al., 2015; Garita-Cambronero et al., 2017; Martins et al., 2020; Merda et al., 2016, 2017; Vauterin et al., 1996). Nonpathogenic xanthomonads have a varied lifestyle with the ability to colonize the plant hosts and survive in various environments outside the plants, such as rain and aerosols (Mechan Llontop et al., 2021; Vauterin et al., 1996). Although referred to as nonpathogenic in the context of their phenotype based on artificial inoculation on the host of isolation, it cannot be ruled out that these strains may cause disease in other hosts. Some of these nonpathogenic *Xanthomonas* strains have been isolated together with pathogenic relatives from a diversity of host plants, at times, from the same lesion in infected plants or asymptomatic hosts or seed-lots or transplants (Gitaitis, 1987; Vauterin et al., 1996). Some of these nonpathogenic xanthomonads are opportunistic pathogens under favorable conditions (Vauterin et al., 1996) and have amylolytic and/or pectolytic activity, which allows them to cause soft rot on their host (Gitaitis, 1987; Zarei et al., 2022). Vauterin et al. (1996) systematically characterized seventy diverse nonpathogenic xanthomonads based on fatty acid methyl ester (FAME) and sodium dodecyl sulfate-polyacrylamide gel electrophoresis (SDS-PAGE) protein patterns. This pioneering study indicated potential new species and the need to address the diversity and relatedness of nonpathogenic strains to the pathogenic strains of xanthomonads from an ecological viewpoint and the practical aspects of disease diagnostics and management strategies. In the last two decades, several studies have addressed this question of diversity, focusing on individual species, using more advanced methods of multi-locus sequence typing and genome sequencing (Bansal et al., 2020, 2021; Cesbron et al., 2015; Essakhi et al., 2015; Gonzalez et al., 2002; T. Li et al., 2020; Triplett et al., 2015). Whole genome-based phylogeny placed the crop-associated nonpathogenic xanthomonads in the species *arboricola*, *cannabis*, belonging to Group 2 (Cesbron et al., 2015; Jacobs et al., 2015; Merda et al., 2016, 2017) and some in the newly described species, such as *X. sontii* (Bansal et al., 2021), belonging to the early branching clade, Group 1. Apart from *X. arboricola* and *X. campestris*, which house both pathogenic and nonpathogenic strains simultaneously isolated from symptomatic hosts (Lee et al., 2020; Martins et al., 2020), other nonpathogenic strains are only distantly related to the co-colonizing pathogenic strains (Vauterin et al., 1996).

These nonpathogenic strains are diverse in their phylogenetic placements and vary in their makeup of type III secretion systems (T3SS) and associated effectors. The T3SS, encoded by the *hrp2* cluster and type III effectors and/or their repertoires, are important determinants of pathogenicity in xanthomonads, and individual effectors or their repertoires have been hypothesized to contribute towards host specificity (Hajri et al., 2009; Jacques et al., 2016; White et al., 2009). Pathogenic xanthomonads belonging to *X. maliensis*, *X. cannabis*, *X. pseudoalbilineans,* and *X. sacchari* are an exception in that they lack a *hrp2* T3SS, but some possess regulators of T3SS (Triplett et al. 2015; Jacobs et al. 2015, Studholme et al. 2011, Pieretti et al. 2015). Knowing the importance of T3SS and associated effectors in the pathogenicity of xanthomonads, it is unsurprising that most nonpathogenic xanthomonads lack T3SS. Such nonpathogenic xanthomonads lacking T3SS, specifically *X. arboricola*, have been previously referred to as commensal xanthomonads. However, Merda et al. (2017) further showed that some commensal strains of *X. arboricola* possess T3SS but contain only 3-4 effectors. Some commensal strains lack *hrp2* cluster but have up to four effectors (Cesbron et al., 2015).

Interestingly, some *X. arboricola* strains have been reported as weak pathogens based on pathogenicity tests showing faint or mild water-soaking symptoms on the host of isolation (Roach et al., 2018; Sawada et al., 2011). Further genome analysis of weakly pathogenic *X. arboricola* strains revealed a limited set of effectors compared to pathogenic strains, in some cases, up to 16 effectors (Roach et al., 2019), suggesting the possibility of these strains to be potentially pathogenic on other hosts. This heterogenic distribution of T3SSs and variable T3E repertoires make nonpathogenic strains a suitable model to study the evolutionary history of the *hrp2* T3SS family, effectors, and associated regulators in xanthomonads (Cesbron et al., 2015; Merda et al., 2017). Merda et al. (2017) inferred ancestral acquisitions of the *hrp2* cluster in *Xanthomonas* and indicated loss or subsequent gain in certain clades in 82 genomes spanning the genus *Xanthomonas*. As we uncover additional diversity in the genus *Xanthomonas*, evolutionary gain and loss of different types of T3SSs, including atypical T3SSs (Pesce et al., 2017; Pieretti et al., 2015) in xanthomonads along with effectors and regulators can be a valuable approach to understand the role of T3SSs in allowing intimate association of xanthomonads with plants and to further determining their lifestyle.

The presence of nonpathogenic xanthomonads in association with pathogenic xanthomonads from the same infected tissues raises the important question of whether the presence of nonpathogenic xanthomonads influences the population dynamics of pathogenic xanthomonads and vice versa, either through the exchange of genetic material or through sharing of public goods, such as cell-wall degrading enzymes or their products of degradation (Sadhukhan et al., 2023). Profiling of the mobile genetic elements (MGEs) shared between pathogenic and nonpathogenic strains can provide critical information to understand how genes and their encoded functions can be exchanged via horizontal gene transfer. These MGEs can influence selection pressure-driven changes in the population dynamics of pathogens and nonpathogens. Such events linked to pathoadaptation have been proposed (Cesbron et al., 2015; Meline et al., 2019).

*Xanthomonas* strains were recently found in the endosphere of *Arabidopsis* as a part of the At-LSPHERE collection. Evaluation of these strains’ pathogenicity on wild-type *Arabidopsis* and immunocompromised plants confirmed their opportunistic or conditional pathogenic nature based on aggressive symptoms on immunocompromised plants lacking plant NADPH oxidase (RBOHD) (Pfeilmeier et al., 2021). This study further highlighted the role of microbial community members and microbiota-induced plant immunity in reducing the prevalence of opportunistic strains. On the other hand, a closely related endophytic *Xanthomonas* strain, WCS2014-23, was identified as a member of the consortium recruited to the rhizosphere of *Arabidopsis thaliana* upon foliar infection with the biotrophic pathogen and played a role in induced systemic resistance against the biotrophic pathogen and enhancing plant growth (Berendsen et al., 2018). These findings raise the question of whether nonpathogenic xanthomonads play an important role as resident and functional members of the phyllosphere microbiome or are transient nonfunctional community members. A related question is if the so far uncharacterized xanthomonads that have been isolated from rainwater are *bona fide* phyllosphere microbiome members that are only transiently present in the atmosphere or if they present a separate population of non-plant-associated xanthomonads (Failor et al., 2017).

In this study, we set out to address the above-mentioned knowledge gaps on the overall diversity of nonpathogenic xanthomonads regarding the genetic diversity of strains, associated virulence factors, and mobile genetic elements. We also focused on understanding the evolution of pathogenic and nonpathogenic strains and deciphering genes associated with the adaptation of these diverse strains to different lifestyles in association with plants and the environment. We sequenced a collection of 83 presumptive nonpathogenic *Xanthomonas* strains from diverse hosts, environments, and geographical locations. The phylogenetic placement of these strains spanned the entire genus, including the identification of potentially new species. The heterogeneous distribution of T3SS and associated effectors across pathogenic, weakly pathogenic, and commensal strains suggested a lack of apparent association of these important pathogenicity factors with the lifestyle of *Xanthomonas* strains. Thus, we used an integrated approach of comparative genomics and association analysis to identify the genomic attributes associated with these lifestyles of strains spanning the genus phylogeny.

## Materials and Methods

### Bacterial strains collection and genome sequencing

Nonpathogenic *Xanthomonas* strains collected from different plant hosts and environmental samples (Table S1) were used for genomic DNA extraction using the CTAB-NaCl method (William et al., 2012). Degradation and contamination of the genomic DNA were monitored on 0.5% agarose gels. DNA concentration was measured using a Qubit® DNA Assay Kit on a Qubit® 2.0 Fluorometer (Life Technologies, CA, USA) and submitted to the Joint Genome Institute (JGI) for library preparation and sequencing. Paired-end reads were generated by multiplexing 12 libraries in a single lane on the Illumina NovaSeq (PE150) platform. Raw reads, annotation data, and final assembly are in the JGI data portal (http://genome.jgi.doe.gov). The information for 83 newly sequenced genomes from this study is in Table S1.

### Genome-based identification of *Xanthomonas* strains

Comparative genomic analysis was performed among 134 *Xanthomonas* strains, including 83 strains from this study and 51 representative *Xanthomonas* strains from NCBI (Table S2). Average nucleotide identity (ANI) was estimated using all-versus-all strategies using FastANI (v1.1) (Jain et al., 2018) and pyani (v0.2.12) (Pritchard et al., 2015). We also used the ANI values from the web server LINbase (Tian et al., 2020) and Microbial Species Identifier (MiSI) (Varghese et al., 2015) as additional tools in species circumscription. The MiSI method addresses inconsistencies based on ANI alone and includes alignment fractions and genome-wide ANI values. Additionally, twenty strains representing the novel species diversity were subject to the Type (Strain) Genome Server (https://tygs.dsmz.de/) to calculate digital DNA-DNA hybridization (dDDH) values based on the Genome BLAST Distance Phylogeny (Meier-Kolthoff et al., 2013). A combination of ANI, dDDH, and MiSI was used to designate the ";new species"; status to a given strain only when values were below the accepted threshold (≤95% for ANI and ≤70% for dDDH) (Kim et al., 2014).

The whole proteome of the 134 *Xanthomonas* strains was compared by OrthoFinder (v2.5.2) (Emms & Kelly, 2019) to identify orthogroups using the original algorithm (Emms & Kelly, 2015). The identified orthogroups were used to infer unrooted gene trees using the BLAST-based hierarchical clustering algorithm DendroBLAST (Kelly & Maini, 2013). Using this set of unrooted gene trees, the STAG algorithm identified the closest pair of genes from those species to infer an unrooted species tree (Emms & Kelly, 2018). The unrooted species tree inferred from STAG was then rooted using the STRIDE algorithm (Emms & Kelly, 2017) by identifying well-supported gene duplication events. The resulting cladogram was visualized with R package *ggtree* (G. Yu et al., 2017).

To identify and visualize possible conflicting signals that would suggest recombination events and evolutionary relationships within the *Xanthomonas* sequence data, Multi-locus sequence analysis (MLSA) was carried out for 12 housekeeping genes fragments (*gyr*B, *gap*A, *lac*F, *glt*A, *fsu*A, *lep*A, *atp*D, *rpo*D, *gln*A, *ef*P, *dna*K, and *fyu*A) using autoMLSA2 (v0.7.1) (https://github.com/davised/automlsa2). The resultant splits.nex file was used in SplitsTree4 (v4.17.0) (Huson et al., 2008) to develop a phylogenetic network. The possibility of recombination events was identified by the branches that form parallelograms (Joseph & Forsythe, 2012).

### Analysis of the gain and loss dynamics of the T3SS clusters

Protein sequences of T3Es representing all effector families, putative effectors (Table S4), and their diversity were also identified in genome sequences using tBLASTn searches. The T3SS-coding genes from five different *Xanthomonas* species (*X. campestris* pv. *vesicatoria* 85-10, *X. campestris* pv. *campestris* ATCC33913, *X. translucens* pv. *translucens* DSM18974, *X. albilineans* CFBP 2523, and *Xanthomonas* sp. 60), were used as query to perform BLASTn searches on *Xanthomonas* strains genomes using autoMLSA2 (v0.7.1) with the cut-offs set to 40% identity and 30% coverage. Heatmaps for the blast searches were generated using the *R* package *Pheatmap* (v1.0.12).

Branch-specific gain and loss probabilities of *Xanthomonas* T3SS genes during the evolution were inferred with the species tree and presence/absence in the 134 genomes using GLOOME (Cohen et al., 2010). GLOOME analyzes presence and absence profiles (phyletic patterns) and accurately infers branch-specific and site-specific gain and loss events. We first inferred the gain and loss dynamics of all genes encoding components of the T3SS. Specifically, we searched the genes of four T3SS clusters: (1) *hrp2* cluster derived from *X. campestris* pv. *Campestris* ATCC33913 (*Xcc* cluster, 22 genes); (2) *X. translucens* pv. *translucens* DSM18974 (*Xtra* cluster, 18 genes); (3) *X. albilineans* CFBP 2523 (*Xalb* cluster, 11 genes); (4) *Xanthomonas.* sp. 60 (strain 60 cluster from this study, 11 genes). We additionally searched for the presence and absence of genes encoding transcription factors involved in T3SS-related pathogenicity: HrpG, HrpX, HpaR, and HpaS (four genes). These 66 genes were searched in the 134 genomes using BLASTp. A hit was considered if the identity percentage was at least 50%, the E-value was lower than 10^-10^, and the coverage was at least 30%. When a hit was detected to two or more genes from different T3SS clusters, the one with the highest bit-score was retained. As the phylogenetic tree for the analysis, we used the species tree generated by OrthoFinder. The final GLOOME analysis was performed with default parameter values. Graphical visualization of the tree was done using FigTree (v1.4.4) (http://tree.bio.ed.ac.uk/software/figtree/).

### Prediction of Mobile Genetic Elements

To study the genomic differences driven by mobile genetic elements (MGE) in the genus *Xanthomonas*, we used Mobile Genetic Element Finder (MGEfinder) (v1.0.6) (Durrant et al., 2020). MGEfinder assembles the short reads and aligns them to a reference genome to find insertions (Durrant et al., 2020). *Xanthomonas* reads from this study and representative short reads were downloaded from NCBI’s SRA database and trimmed using Trim Galore (v0.6.6) (https://github.com/FelixKrueger/TrimGalore). Reference genomes were indexed, and the cleaned reads were aligned with BWA-MEM (v0.7.17) (H. Li & Durbin, 2009). Target strains were assigned to pathogenic reference genomes according to their phylogenetic placement generating eight clusters (Table S2). Each cluster contains a representative/type strain and the strains from this study. The predicted mobile genetic islands were annotated using a consensus from BLASTx results in NCBI, JGI, and UniProt. Islands containing carrier genes of potential interest were further analyzed using JGI BLASTp and gene neighborhood viewer to locate transposases or phage-related genes associated with or flanking the island. Additionally, EasyFig (v2.2.2) (Sullivan et al., 2011) was used to visualize the insertion location of mobile genetic elements within genomes.

### Comparison of secreted carbohydrate-active enzymes

We screened the genomes of commensal, weakly pathogenic, and pathogenic *Xanthomonas* strains for the presence of various genes involved in breakdowns (CEs, PLs, GHs) and assembly (GTs) of carbohydrates, lignin degradation (AAs), and the carbohydrate-binding module (CBM) (Kaoutari et al., 2013; Lairson et al., 2008). Carbohydrate active enzymes (CAZymes) were assigned to Prokka (v1.14.5) (Seemann, 2014) protein output files (.faa files) using run_dbcan command (https://github.com/linnabrown/run_dbcan) against the HMMER, DIAMOND, and eCAMI databases with default settings. Final CAZyme domain annotations were the best hits based on the outputs of at least two databases to investigate the genomic potential of various species for carbohydrate utilization. To assess the impacts of different lifestyles on the secreted CAZyme count while considering phylogenetic signals, pairwise phylogenetic distances were created between the genomes using the function tree.distance() from the package biopython phylo (Cock et al., 2009), which was then used to build a principal component analysis (PCA). The CAZyme count for each genome from the run_dbcan step was then converted into a distance matrix with the function vegdist (method=";jaccard";) from the *Vegan* (v2.6-4) R package (Dixon, 2003) (Oksanen et al. 2009). Permutational multivariate analysis of variance (PERMANOVA) was performed to determine the effect of phylogeny and microbial lifestyle on the distribution of CAZymes with the function adonis2 from the *Vegan* R package. Pairwise comparisons between the lifestyles were carried out using the function *pairwise.perm.manova* (from R package *RVaidememoire*) to understand the difference between lifestyles in terms of their genome content as described in Miyauchi et al. (2020).

### Pathogenicity assays and co-inoculation experiments

A subset of *Xanthomonas* strains isolated from tomato were further tested for their pathogenicity by dip-inoculating tomato plants (susceptible cultivar FL 47R). 4-5 weeks old tomato plants were dip-inoculated with an inoculum of overnight grown *Xanthomonas* strains adjusted to 10^6^ CFU/ml in MgSO_4_ buffer amended with 0.0025% (vol/vol) Silwet L-77 (PhytoTechnology Laboratories, Shawnee Mission, KS, USA) and maintained under greenhouse conditions. The symptoms on leaves and *in-planta* bacterial population were recorded after ten days. The *in-planta* bacterial population was estimated by sampling 2cm^2^ leaf tissue using a cork-borer, homogenizing the tissue in 1ml of MgSO_4_ buffer, and plating on Nutrient Agar using a spiral plater (Neu-tecGroup Inc., NY). Plates were incubated at 28°C for three days, and the population for each *Xanthomonas* strain was determined as colony-forming units per centimeter squared of leaf area.

A coinfection experiment was conducted to gain experimental evidence for the exchange of genetic material among commensal and pathogenic *Xanthomonas* under host selection pressure. Pathogenic *Xanthomonas*, *X. euvesicatoria* 85-10 carrying the intact *avrBs1* gene and the commensal *Xanthomonas* strain T55 were co-inoculated onto pepper cv. Early Cal Wonder (ECW, susceptible cultivar) and ECW-10R (carrying the *Bs1* resistance gene, resistant cultivar) using the dip-inoculation method described above at the inoculum concentration of 10^7^ CFU/ml. The plants were maintained under greenhouse conditions for two weeks, and the leaf tissue containing lesions was sampled from resistant and susceptible cultivars. Inactivation of the *avrBs1* gene by transposons was tested by PCR using primers flanking transposons on transconjugants obtained from these samples.

### Identification of lifestyle-associated genes

We retrieved 1,834 *Xanthomonas* genomes from the NCBI GenBank RefSeq database to ensure a high-quality and minimally biased set of genomes. These genomes were then de-replicated using dRep (v3.2.2) (Olm et al., 2017) to de-replicate the complete dataset using a 95% minimum genome completeness cut-off and 5% maximum contamination. The ANI threshold to form primary clusters (-pa) was set at 0.95 (species level) and 0.99 (strain level) for the secondary cluster. During secondary comparisons, a minimum level of overlap between genomes was set to 80% coverage. The de-replicated genomes were manually curated to include the 83 new genomes generated in this work in addition to representative genomes from diverse *Xanthomonas* species belonging to groups 1 and 2. A total of 337 genomes were selected for downstream analysis (Table S3). OrthoFinder was used as a clustering approach to compare the whole proteome of the 337 *Xanthomonas* strains with the default settings as previously described. To determine significantly enriched or depleted protein clusters in different *Xanthomonas* lifestyles, we used the hypergeometric test, PhyloGLM, and Scoary as described in Levy et al. (2018). Among the three methods, the hypergeometric test looks for the overall enrichment of genes without considering the dataset’s phylogenetic structure. PhyloGLM is a phylogenetic-aware method that eliminates enrichments related to shared ancestry (Ives & Garland, 2010), while Scoary combines the phylogeny-aware test, Fisher’s exact test, and empirical label-switching permutation analysis (Brynildsrud et al., 2016). All these approaches were used on gene presence/absence and gene copy-number data and used for PhyloGLM test. A gene was considered significant a) if it had a q-value < 0.05 for Fisher’s exact test and an empirical *p-*value < 0.05 for Scoary; b) if it had a corrected *p*-value with FDR with q < 0.01 for hypergeometric test; and c) a *p*-value < 0.01 along with an estimate of < −1.5 or > 1.5 in copy number analysis for PhyloGLM. We used eggNOG-mapper (v2.1.7) (Cantalapiedra et al., 2021) to address COG categories to each significantly enriched and depleted protein cluster from the combination of two or more methods (hypergeometric, PhyloGLM, and Scoary) in the different *Xanthomonas* lifestyles. In addition, each ortholog id was queried across the IMG database to obtain annotation based on COG, KO, TIGRFAM, and Pfam. Heatmaps were generated using the *R* package ggplot2.

## Results

### General features of *Xanthomonas* strains from this study, potential new species, and core genome phylogeny

Presumptive nonpathogenic *Xanthomonas* strains sequenced in this study were isolated from various symptomatic and asymptomatic crops, including tomato (*Solanum lycopersicum*) (14 strains), pepper (*Capsicum annuum*) (8 strains), and common bean (*Phaseolus vulgaris*) (27 strains), along with other plant species such as radish (*Raphanus sativus*), walnut (*Juglans*), orange (*Citrus sinensis)*, and sunflower (*Helianthus*). In addition to the strains isolated from plant hosts, 18 strains were recovered from rainwater (Table S1). The genome sizes among the sequenced strains varied from 3.6 Mb for strain 60 to 5.3 Mb for strain F5. The percent GC content ranged from 64.60% for strain 3075 to 69.31% for strain F10. Furthermore, the number of coding sequences (CDS) varied from 3,223 in strain 60 to 4,535 in strain F5. There was no apparent correlation between genome size, CDS, %GC and status as pathogenic and nonpathogenic (Figure S1). While a median genome size of 4.87 Mb and a median number of genes of 4208 are found in the genus *Xanthomonas*, one strain (60) showed a reduced genome size of 3.6 Mb and a reduced number of 3223 coding genes (Table S1).

To determine the taxonomic placement of the 83 newly sequenced strains, ANI, dDDH, and MiSI were used. ANI values of the strains varied from 79% to 100% compared to the representative *Xanthomonas* strains. Nonpathogenic strains sequenced in this study belonged to both *Xanthomonas* groups. For group 1, nine strains belonged to *X. euroxanthea*. For group 2, 34 strains belonged to *X. arboricola*, 16 to *X. cannabis*, 9 to *X. campestris*, and two to *X. euvesicatoria* (Table S5, S6, S7, and Figure S2, S3). The remaining thirteen strains showed ANI values between 85-94% when compared with known species of *Xanthomonas*. These strains were assigned to eight cluster-type cliques or singletons according to the MiSI method, indicating the presence of at least eight potentially novel species (Table S8). Given the findings of potentially novel species adding diversity to the existing *Xanthomonas* genus phylogeny, we established a robust phylogenetic tree based on the single-copy genes of these newly sequenced strains along with type or representative strains of the *Xanthomonas* genus (Figure 1). The OrthoFinder analysis assigned most genes (544,723; 99% of the total) to 11,456 orthogroups. There were 1005 orthogroups in all species, and 819 of these consisted entirely of single-copy genes. The phylogenetic reconstruction showed a considerable diversity of strains isolated from both plant hosts and the environment, and these strains were broadly distributed throughout the genus (Figure 1).

**Figure 1:**
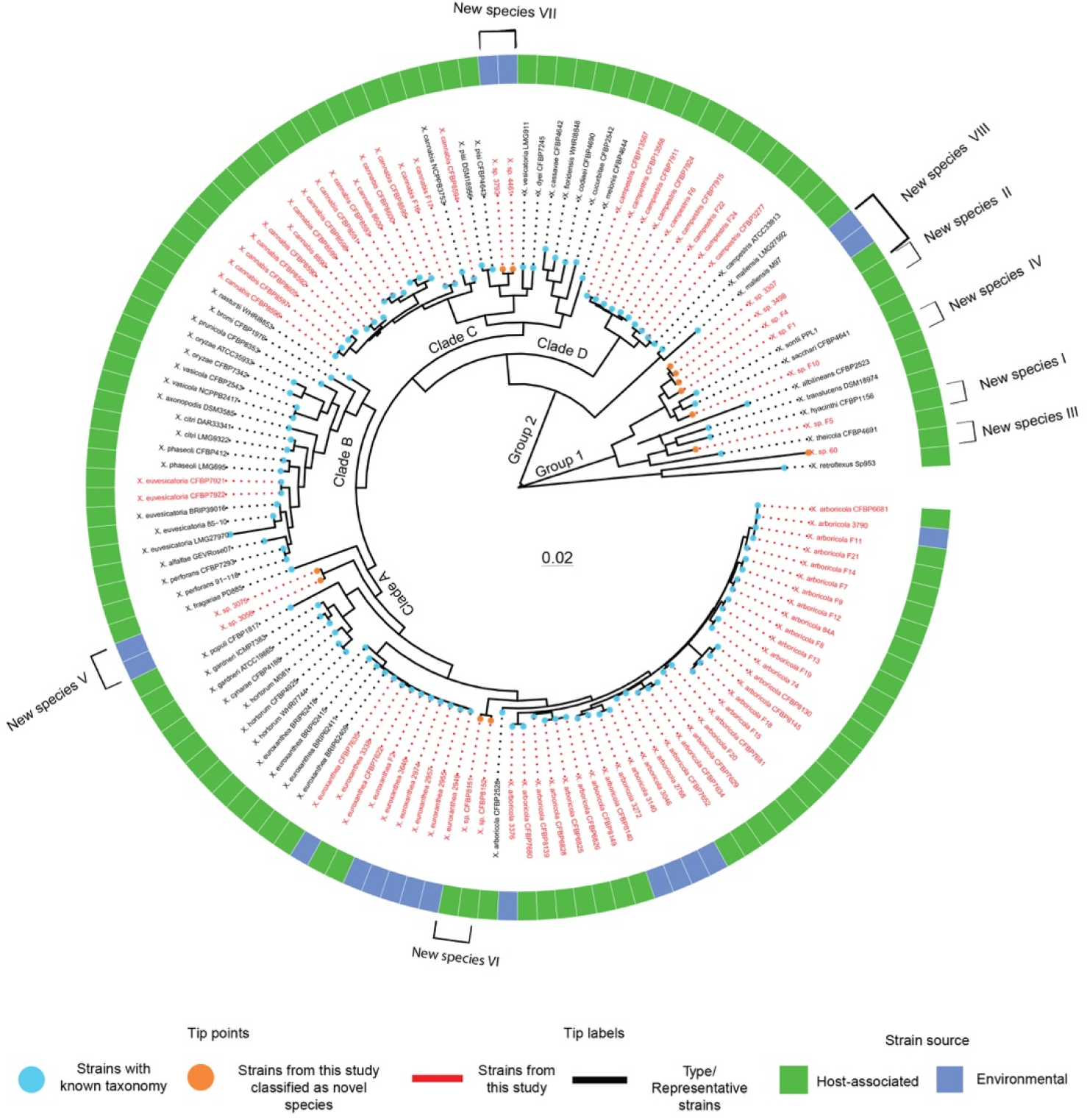
Comparative genome analysis demonstrated the presence of eight novel species in the genus *Xanthomonas*. Maximum-likelihood phylogeny based on the 1005 orthogroups of 134 strains representing the entire *Xanthomonas* genus. The phylogenetic tree was inferred using OrthoFinder v2.5.2 and drawn with *R* package *ggtree*. Here, *Xanthomonas* strains from this study are highlighted in red, while the representative/type strains are in black. The tip points are colored orange to show novel species identified from this study, while cyan represents *Xanthomonas* species with known taxonomy. Blue-colored blocks indicate environmental strains and host-associated strains are indicated by green-colored blocks surrounding the phylogenetic tree.

*X. sontii*, *X. sacchari*, *X. albilineans*, *X. hyacinthi*, *X. translucens*, *X. theicola,* and the two recently described species *X. bonasiae* and *X. youngii* were the only known species belonging to *Xanthomonas* group 1 (Bansal et al., 2021; Mafakheri et al., 2022; Rodriguez-R et al., 2012). The collection sequenced here adds four new *Xanthomonas* species to group 1, species I (strain F5), species II (strain F1), species IV (strain F10), and species VIII (strains 3307, 3498, and F4), all isolated from citrus plants and rainwater. Another new *Xanthomonas* species, species III (strain 60), from this collection clustered with early branching species at the base of the phylogenetic tree, along with *X. retroflexus*.

Our collection also added three new species to *Xanthomonas* group 2 (Figure 1). Most strains sequenced here belong to clade A, specifically to *X. arboricola* (Table S5, S6) and *X. euroxanthea* (Table S7). Strains isolated from bean seeds (CFBP 8151 and CFBP 8152) were closely related to *X. arboricola,* and strains isolated from rainwater (3075 and 3058) belong to potentially novel species group, species VI and species V, respectively, within clade A (Table S8, Figure 1). Two of our sequenced strains belonged to clade B and were identified as *X. euvesicatoria* (CFBP 7921 and CFBP 7922). Strains in clade C isolated from bean seed, tomato, and nightshade plants belong to *X. cannabis* species (16 strains). This clade also harbors one novel *Xanthomonas* species, species VII (strains 3793 and 4461), isolated from rainwater. Crop-associated *Xanthomonas campestris* strains (9 strains) isolated from radish, bean, and tomato plants belong to clade D, including pathogenic *X. campestris* strains (Figure 1).

### Distribution of T3SS clusters across the phylogeny

We screened three types of T3SS clusters known within the genus *Xanthomonas* across the set of genomes: (1) the *hrp2* cluster present in group 2 xanthomonads (Tampakaki et al., 2010), (2) the SPI-1 type present in *X. albilineans* (Pieretti et al., 2009), and (3) the noncanonical T3SS cluster present in *X. translucens* (Wichmann et al., 2013). In addition, we also included the T3SS cluster from *Stenotrophomonas* sp. to represent the *sct*-type cluster from a closely related genus. Among the 83 newly sequenced *Xanthomonas* genomes, 24% contained a functional T3SS cluster (Figure S4). Strain *Xanthomonas* sp. 60 showed the presence of a unique *sct*-type T3SS cluster with a gene organization comparable to the one found in *Stenotrophomonas chelatiphaga* DSM 21508 (Figure S5). Apart from partial T3SS clusters in strains of *X. cucurbitae and X. fragariae*, we also identified single gene encoding protein V of sct-type cluster in *X. cannabis* CFB P8595, *X. cannabis* 8600, *X. cannabis* CFBP 8600, *X. arboricola* F12, *X. arboricola* 84A, and *Xanthomonas* sp. 4461 and a single *hrpF* in *X. arboricola* CFBP 7681 (Figure 2). T3SS clusters were missing in *X. pisi*, *X. floridensis*, *X. melonis*, *X. maliensis*, *X. sontii*, *X. sacchari*, and *X. retroflexus. X. phaseoli* pv. *phaseoli* CFBP 412 showed the presence of two types of T3SS, the *hrp2* cluster and the SPI-1 type cluster present in *X. albilineans*, like previous findings on *X. phaseoli* pv. *phaseoli* CFBP 6164 (Alavi et al., 2008) (Figure 2).

**Figure 2:**
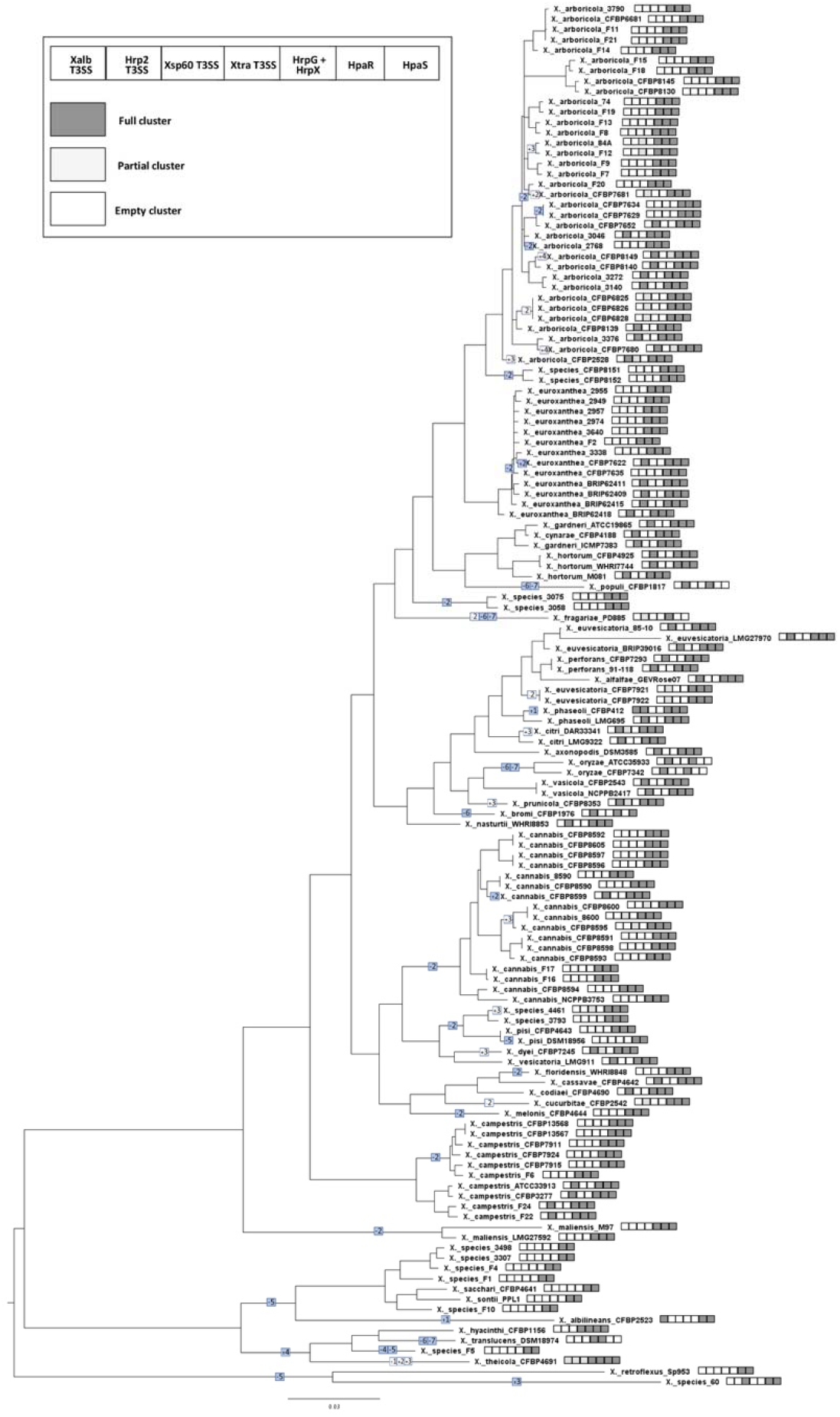
Multiple events of gain and loss of T3SS were evident in the genus *Xanthomonas* along with the presence of a novel T3SS cluster. Maximum-likelihood phylogeny based on the core proteome and T3SS cluster and regulators gain-loss prediction inferred for the 134 strains representing the entire *Xanthomonas* genus. The presence/absence of the different types of T3SS and regulators are represented by an ordered vector of size 7, such that a dark grey, a light grey, and a white i^th^ element in the vector indicates a full presence, a partial presence, and an absence of the i^th^ element, respectively. Inferred full and partial T3SS clusters are in color or white background, respectively. Acquisition and loss events are represented by plus and minus signs, respectively. For example, a colored +2 indicates a full acquisition of the 2^nd^ T3SS in this figure, i.e., *hrp2*.

### T3SS was gained and lost multiple times in the genus *Xanthomonas*

To better understand the evolution of T3SS clusters in the genus *Xanthomonas*, we next studied the presence and absence patterns of T3SS clusters among the analyzed genomes. The phyletic pattern generated above was input to GLOOME, which maps gain and loss events onto the phylogeny. According to the scenario estimated with GLOOME, there were several independent acquisition and loss events of T3SS clusters during the evolution of xanthomonads. Independent acquisition of the *sct*-type T3SS cluster was inferred to have occurred in *Xanthomonas* sp. 60 (Figure 2). The ancestor of *X. translucens, X. hyacinthi, Xanthomonas* sp*. F5,* and *X. theicola* acquired the *Xtr*-type T3SS cluster. This cluster was found to be conserved in the *X. translucens* clade. However, it has been subsequently lost in *Xanthomonas* sp. F5 and partially lost in *X. hyacinthi* and *X. theicola,* as indicated by a partial *Xtr*-type cluster (Figure S5 and Figure 2). The SPI-1 T3SS cluster was independently acquired in *X. albilineans* and *X. phaseoli* CFBP 412.

Next, in addition to the probabilities of gene-gain/loss estimated by GLOOME analysis, we noted the genomic context and sequence identities to identify *hrp2* cluster gain/loss events associated with group 2 xanthomonads. Based on the sequence identities of *hrp2* cluster genes, clades A, C, and D were observed to possess the *Xcc*-type *hrp2* cluster, while clade B, except for *X. nasturtii*, possessed the *Xeu*-type *hrp2* cluster. It is possible that replacement of the *Xcc*-type *hrp2* cluster by the *Xeu*-type *hrp2* cluster occurred within clade B strains, except in *X. nasturtii*, through rearrangements.

According to GLOOME analysis, regulators of T3SS, HrpX, and HrpG, were acquired by a common ancestor of *Xanthomonas* before the split into group 1 and group 2. A single loss event of these regulators occurred in group 2, where these genes were lost on the branch leading to *X. pisi* DSM18956. In group 1 these genes were lost in several independent events: (1) on the branch leading to the common ancestor of *X. albilineans*, *X. sontii*, *X. sacchari*, and *Xanthomonas* sp. strains (F10, F1, F4, 3307, and 3498); (2) on the branch leading to *Xanthomonas* sp. F5; (3) on the branch leading to the common ancestor of *X. retroflexus* and *Xanthomonas* sp. 60. An alternative less parsimonious explanation is that these regulators were only acquired by group 2 *Xanthomonas* and were independently acquired by the cluster of *X. hyacinthi*, *X. translucens*, *X*. *theicola, and Xanthomonas* sp. F5, followed by loss of these genes together with loss of *Xtra*-type T3SS genes in *Xanthomonas* sp. F5. Similar to HrpX and HrpG, the regulators HpaR and HpaS were also acquired by the common ancestor of *Xanthomonas* before the split of group 1 and group 2. These regulators were lost independently on numerous occasions: (1) on the branch leading to *X. translucens* in group 1; (2) on the branch leading to *X. populi* in group 2, and (3) on the branch leading to *X. oryzae* in group 2.

### T3E repertoire ranges from zero to forty-one in crop-associated and environmental strains

Previous studies involving genome screening for T3E of nonpathogenic *X. arboricola* strains indicated low effector gene loads, with a reduced core effector set ranged from zero to four effectors, namely, XopR, HpaA, XopF1, and AvrBs2 (Merda et al., 2017). Given the diversity of commensal xanthomonads spanning the entire phylogeny of the genus *Xanthomonas* and the fact that this study also included environmental strains, we hypothesized that low effector loads might be widespread among nonpathogenic xanthomonads that are either plant-associated or environmental strains, owing to their global broad host range. However, we caution that host range tests have not been conducted for each strain sequenced in this study. Thus, we refer to a global broad host range based on previous studies that either recovered nonpathogenic isolates from diverse plant hosts or tested their host range on diverse plants. Our analysis indicated that effector repertoires vary greatly from zero to 41 (Figure S6).

*X. arboricola* strains sequenced in this study, although belonging to a monophyletic group, showed a considerable variation in the presence/absence of T3SS and the size of effector repertoires, ranging from one to twelve known effectors. Strains lacking T3SS but containing effectors, XopAW and XopAX, included crop-associated and environmental strains. Some crop-associated strains lacking a T3SS possessed an additional T3E, AvrBs2. Two *X. arboricola* strains (F21 and CFBP 6681) isolated from tomato lacked T3SS but possessed two effectors, AvrBs1 and XopH, in addition to XopAW and XopAX. AvrBs1 and XopH have been identified as plasmid-borne effectors in *X. euvesicatoria*, a tomato pathogen. Interestingly, a set of strains isolated from rainwater and from diverse crops such as walnut, pepper, and bean seed shared the same effector repertoire (XopR, XopF1, XopF2, AvrBs2, and XopAW), in addition to a functional T3SS. Strains of *X. arboricola* (CFBP 6825, CFBP 6826, and CFBP 6828) isolated from pepper possessed unusually large effector repertoires comprising 10 T3Es (XopZ2, XopR, XopP, XopF1, XopF2, XopAW, XopAR, XopAL1, XopAD, AvrBs2) comparable to those found in *X. arboricola* strains pathogenic on walnut (Figure S6).

We hypothesized that *X. arboricola* strains with intact T3SSs and larger effector repertoires (i.e.,> seven effectors) would be weakly pathogenic on their host of isolation. Pathogenicity assays using the dip-inoculation method mimicking a natural infection and *in-planta* population growth were performed using six *X. arboricola* strains (CFBP 6825, CFBP 6826, CFBP 6828, CFBP 6681, CFBP 7681, and CFBP 7680) possessing different repertoires of T3Es and presence/absence of a T3SS in tomato cv. FL47R. Strains CFBP 6825, CFBP 6826, and CFBP 6828 triggered slight disease symptoms on tomato leaves at 10 days after inoculation (DAI) (Figure S7A). Strains CFBP 6681 and CFBP 7681 were unable to colonize the host tissue, while strains CFBP 6825, CFBP 6826, and CFBP 6828 maintained a population of ∼ 10^5^ CFU/cm^2^ at 10 DAI (Figure S7B). No visible symptoms were observed for strain CFBP 7680, despite maintaining a population of ∼ 10^4^ CFU/cm^2^ at 10 DAI (Figure S7B). A comparison of the T3Es among these strains indicated that strains that caused disease symptoms had a repertoire of T3E of similar size (11 effectors). In contrast, strains that were unable to cause disease lacked T3SS and had few effectors (<7 effectors) (Figure S7C).

Along with the variable presence of T3SSs, crop-associated nonpathogenic strains also varied in effector repertoire size, which ranged from one to 10 effectors in *X. cannabis* and reached 25 effectors in *X. campestris* (Figure S6). These strains with a higher number of effectors (>7) may suggest their possible pathogenic status, although their host range needs further exploration. Rain-derived *X. euroxanthea* strains lacked T3SSs and possessed a single effector, XopR. Crop-associated *X. euroxanthea* strains isolated from tomato and bean contained a T3SS and effectors, XopF1, XopF2, XopZ2, XopAK, and XopR. Rain-derived novel *Xanthomonas* sp. strains (3058, 3075, 3793, and 4461), although lacking canonical T3SSs, possessed orthologs of the HrpG/X master regulators. Strains 3058 and 3075 lacked any known effectors, while strains 3793 and 4461 contained homologs of AvrXccA1 and AvrXccA2. XopAW and AvrXccA1 were observed in the crop-associated strains 3793 and 4461 lacking T3SSs but containing sequences homologous of HrpG/X.

Based on the work described so far, we defined the lifestyle of strains based on the presence of the T3SS gene cluster and putative effectors. Strains that possessed an intact T3SS and 13 or more T3Es were considered as potentially pathogenic strains. *Xanthomonas* strains with a number of T3SS effectors ranging from 7 to 12 were considered potentially weak pathogens, while strains lacking an intact T3SS and having <7 or no T3Es were considered commensals (Table S3). These three types of lifestyles defined here will be used for the following downstream analyses to infer genetic exchange between commensal and pathogenic *Xanthomonas* and to identify the features that define these lifestyles.

### Genetic exchange between commensal, weakly pathogenic, and pathogenic *Xanthomonas* **strains**

To determine whether environmental (i.e., rainwater-associated) or crop-associated commensals or weakly pathogenic strains exchange genetic material with crop-associated pathogenic strains, we examined phylogenetic networks inferred from concatenated sequences of twelve housekeeping genes using SplitsTree (Figure 3). This analysis revealed reticulated events between commensal and pathogenic *Xanthomonas* strains, suggesting recombination. Two evident reticulation events were identified at the intersections of the network, one between pathogenic and commensal or weakly pathogenic strains belonging to *X. arboricola*, and another one between species encompassing clades B, C, and D of group 2 and species belonging to group 1, indicating the flow of genetic information between them (highlighted gray in Figure 3). Three intrinsic events can be observed by closely analyzing the parallelograms between Group 1 and Group 2. The first event is localized in the central part of the entire network. This reticulated event links the two main branches involving all species belonging to clades B, C, and D of group 2 and species belonging to group 1 (highlighted purple in Figure 3). The second event involves clade D (*X. campestris*), some species belonging to clade C and the entire group 1 (highlighted orange in Figure 3). Finally, the third event is exclusively shared between species from clade B and some species belonging to clade C (highlighted green in Figure 3). According to the neighbor-net tree, at least two strains isolated from rainwater appear to result from the above-mentioned putative recombination events. Within the clade A parallelogram, *X. arboricola* strains (3790, 2768, 3140, 3272, 3046, and 3376) isolated from rainwater showed genetic recombination with type strain *X. arboricola* pv. *juglandis* CFBP 2528 and other crop-associated commensals/weakly pathogenic *X. arboricola* strains. A similar observation was found for *Xanthomonas* sp. strains (3793, 4461, 3307, and 3498) as these environmental strains exchanged genomic content with the ancestors of crop-associated pathogenic and commensal species belonging to clade C and of species belonging to Group 1, respectively.

**Figure 3:**
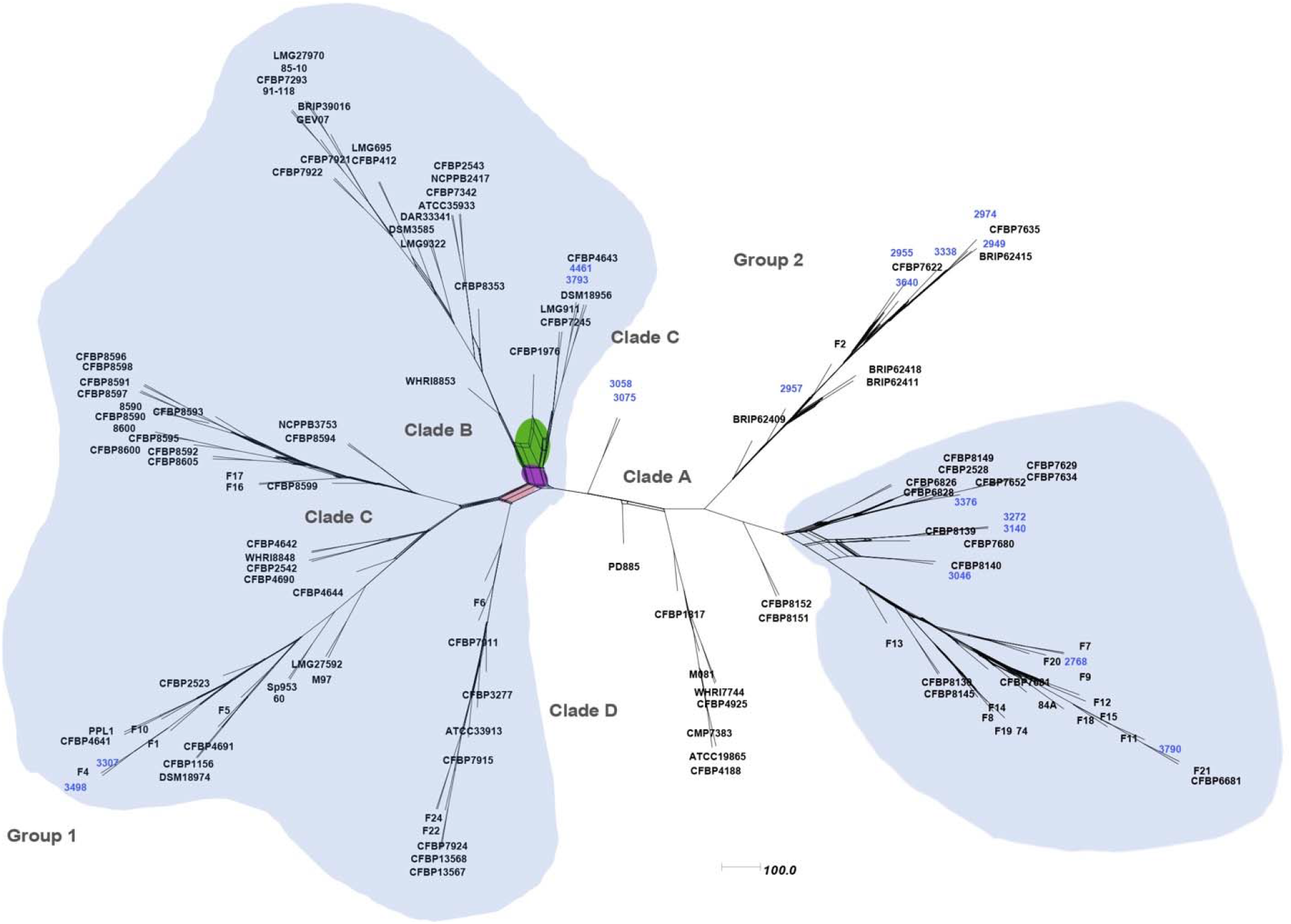
Phylogenetic network between pathogens, weak pathogens, and commensals suggests the possibility of several recombination events during their evolutionary history. Neighbor-net tree constructed using SplitsTree software based on concatenated 12 housekeeping gene sequences generated for the 134 *Xanthomonas* strains, indicating diversity and recombination events.

Many exchange events were observed between the ancestors of crop-associated commensal, weakly pathogenic, and pathogenic strains isolated from the same crop hosts. Within the *X. arboricola* clade, two commensal strains (CFBP 7629 and CFBP 7634) and one weakly pathogenic strain (CFBP 7652) isolated from walnut and the pv. *juglandis* pathotype strain (CFBP 2528) showed a reticulated network. *X. campestris* ATCC33913, pathogenic on crucifers also appears to result from the genomic exchange with the ancestor of the commensal strains *X. campestris* (CFBP 13567 and CFBP 13568), both isolated from radish plants.

The sympatric association of commensals with pathogens, the presence of shared T3Es in commensals, and the observation of recombination signals between commensals and pathogens suggest that commensals and pathogens may act as repositories of fitness traits for each other. To test this hypothesis, we co-inoculated *X. cannabis* T55, a nonpathogenic strain with the T3E gene *avrBs1* with a transposon insertion, IS476, and a plant pathogenic strain, *X. euvesicatoria* 85-10, on pepper plants. *X. euvesicatoria* 85-10 harbors a functional *avrBs1* gene. AvrBs1 induces a hypersensitive response (HR) when infiltrated in the resistant pepper genotype Early California Wonder 10R containing the *Bs1* resistance gene. When screening for putative *X. euvesicatoria* transconjugants, we found that none of the strains isolated from the susceptible pepper genotype (Early California Wonder) had acquired the disrupted avirulence gene from strain T55. In contrast, of nine pools each containing 50 transconjugants, at least two pools isolated from the resistant pepper genotype tested positive for an IS476 transposon insertion (Figure S8).

By screening for mobile genetic elements and associated genes using a computational approach, we further tested the hypothesis that commensal xanthomonads may act as reservoirs carrying fitness and virulence factors that can potentially be transferred to other strains through mobile genetic elements. MGEfinder identified at least one predicted mobile genetic island for each phylogenetic cluster and over 300 unique MGEs. Predicted mobile islands containing genes of possible interest and all islands identified as having a mobility gene, such as transposases or integrases, are listed in Table S9A. While many predicted mobile genetic elements that we found contain housekeeping genes, some contained genes that may play a role in increasing fitness or virulence. The predicted mobile islands contained genes associated with antimicrobial resistance and genes for bacterial and fungal competition, such as multi-drug efflux pumps, type IV secretion system genes, chitinase, and virulence factors. Most mobile genetic elements also contained integrase and phage-related genes. Type III effector XopAD, flanked on both ends by IS5 transposases, was observed in *X. campestris* strains F24 and F22. This island was also found in *X. campestris* pv*. raphani* 756C (the insertion location is shown in Figure S9). The type III effector XopAA, flanked by a putative transposase, was found in *X. euvesicatoria* strains CFBP 7922 and CFBP 7921. An endopolygalacturonase, known for degrading pectin in plant cell walls (Federici et al., 2001), was identified in *Xanthomonas* sp. F1. A peptidoglycan O-acetylase, which may alter bacterial cell walls to avoid lysis by an innate immune response (Sychantha et al., 2018), was detected in *X. arboricola* 3272 (Table S9B).

### Lifestyle had a significant effect in determining the repertoires of cell wall-degrading enzymes in *Xanthomonas*

We hypothesized that nonpathogenic *Xanthomonas* strains possess a distinct repertoire of cell-wall degrading enzymes compared to that from pathogens. Such repertoire might confer them the ability to utilize a wide range of carbohydrate substrates and colonize diverse host plants. Within the genus *Xanthomonas*, different species generally had similar types of CAZymes, but with large variations in the absolute numbers of genes within each category in the CAZy profiles (Figure 4, Table S10). We assessed the effect of bacterial lifestyle on repertoires of genes coding for CAZymes. Distance-based redundancy analyses (dbRDA) of Jaccard distances and permutational multivariate analysis of variance (PERMANOVA) calculated on the genomic compositions of CAZyme family revealed a significant contribution of lifestyle to the distribution of gene repertoires for CAZyme (R^2^ = 0.06, *p*□<□0.05). Furthermore, a pairwise comparison of genomes of different lifestyles revealed CAZyme gene repertoire composition across commensals, weak pathogens, and pathogens to be significantly different (*p* < 0.05), suggesting that lifestyle plays an important role in determining the distribution of CAZymes. Principal component analysis on a matrix containing CAZymes using Jaccard distances showed increased separation of pathogenic from weakly pathogenic strains and commensals.

**Figure 4:**
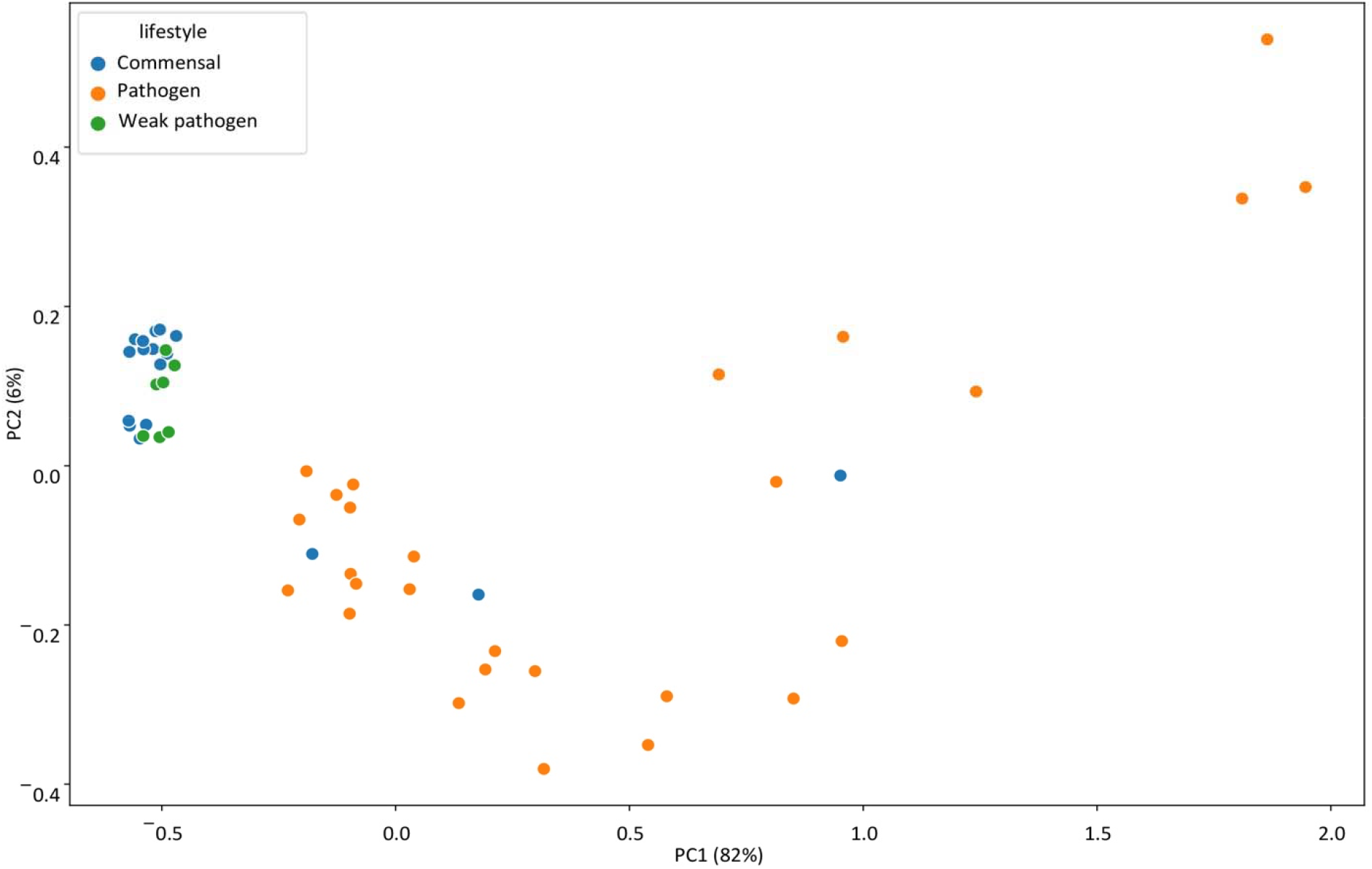
*Xanthomonas* lifestyle can be explained by altered CAZymes landscape. Principal Component Analysis (PCA) plot showing the contribution of the different bacterial lifestyle in the distribution of gene repertoire for carbohydrate active enzymes (CAZyme).

### Association analysis to identify genes and features that define the lifestyle of *Xanthomonas* **species**

Next, we were interested in identifying genes either involved in the adaptation of *Xanthomonas* species to a commensal or a pathogenic lifestyle. Association analysis was performed on the 26,812 orthogroups from a set of 337 *Xanthomonas* genomes containing representative strains from different clades that comprised both pathogens, commensals, and weak pathogens (Table S4, Figure S10). We hypothesized that commensals or weak pathogens and pathogens possess genes or gene families that define their adaptation to the respective lifestyles and have evolved these genes in a phylogeny-independent manner. Presence/absence as well as copy number matrix of orthologs, were used as input to conduct association analysis using three methods, Scoary, PhyloGLM, and hyperglm. Candidate genes present/absent or enriched/depleted in commensal, weakly pathogenic, and pathogenic strains were identified (Figure S11). Overall, each lifestyle category had unique genomic attributes. Interestingly, weak pathogens, as defined here by the presence of a T3SS and 7-12 effectors, contained overlapping genes with both pathogens and commensals (Figure S11).

Among the genes identified as enriched in commensals compared to pathogens and weak pathogens, we found those that belonged to the following functional categories: intracellular trafficking and secretion (type VI secretion system proteins and putative effectors), carbohydrate metabolism and transport, phages and transposons, post-translational modification, protein turnover and chaperones, replication and repair and defense mechanisms (specifically those encoding multi-drug efflux proteins) (Supplementary table S11, Figure 5). Enrichment of genes involved in carbohydrate metabolism in commensals may indicate their ability to utilize broader energy sources than pathogens. A distinct set of DNA-binding transcriptional regulators in commensals may allow them to easily switch between different energy sources depending on their availability on a diverse group of hosts. Other genes that may also impart stress tolerance related to a much broader range of conditions associated with plants or environments outside plants include multi-drug efflux proteins, chaperones, outer-membrane transporters, and type I restriction-modification system genes. During epiphytic colonization of a broad range of hosts, enrichment of genes involved in type VI secretion systems and effectors may allow the commensal xanthomonads to mediate interactions with the resident epiphytic community or pathogenic species (Drebes Dörr & Blokesch, 2018).

**Figure 5:**
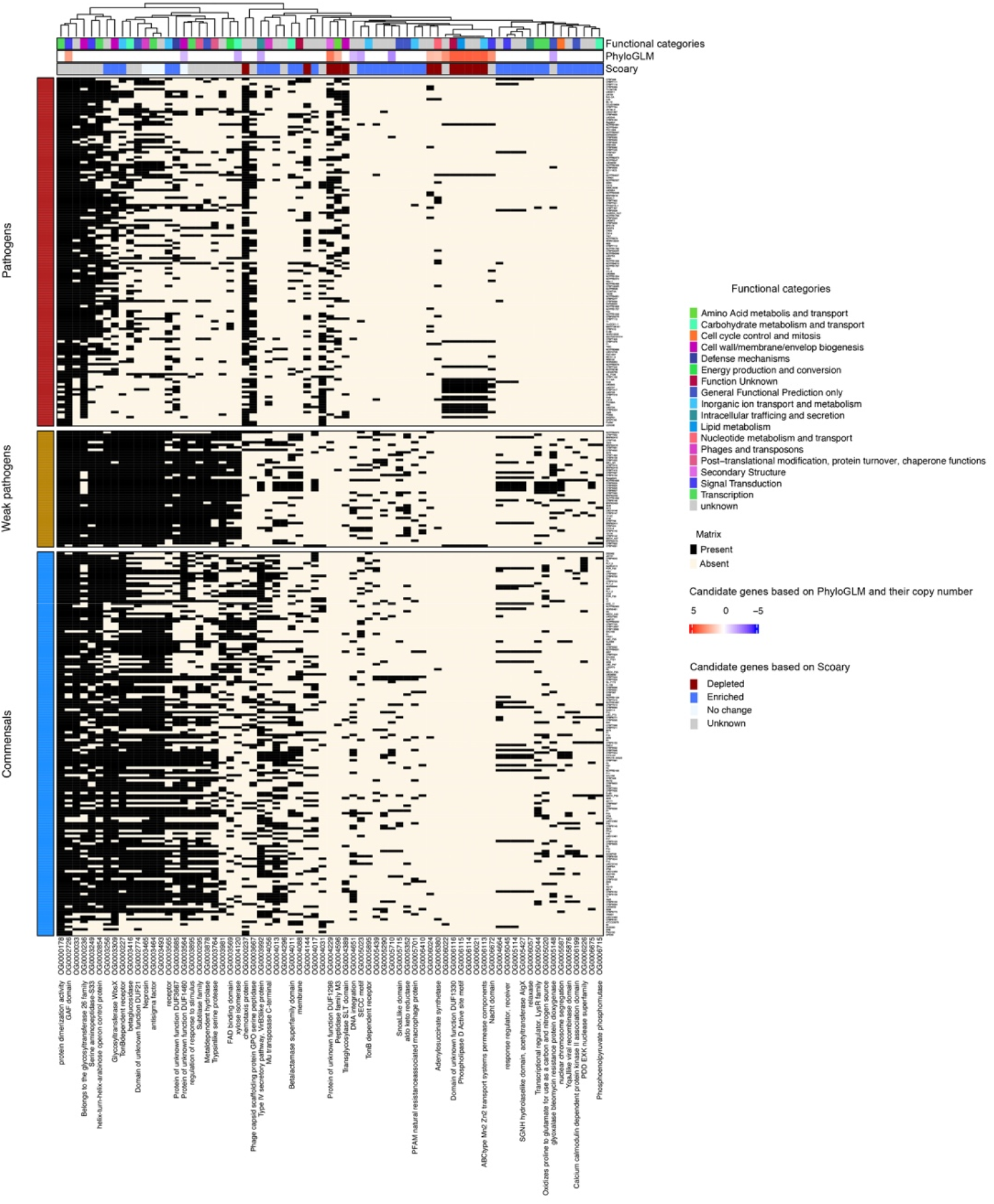
Genomic architecture of *Xanthomonas* contains signatures for both phylogenetic placement and their associated lifestyle. A complex heatmap shows the select candidate genes associated with commensal, weakly pathogenic, and pathogenic lifestyles. The candidates obtained from different methods in figure S11 were further narrowed by identifying those present/enriched in commensals and weak pathogens compared to pathogens and vice versa. The matrix shows the presence/absence of genes across these lifestyles along with their functional categories and annotations on the Y-axis.

We next identified the genomic attributes shared by commensals and weak pathogens but absent from pathogens that could explain strategies evolved by nonpathogens during their association with a broad range of hosts (Table S12A, S12B, Figure 5). Glyoxalase bleomycin resistance protein dioxygenase was identified to be enriched in both the commensal and weakly pathogenic xanthomonads. This protein might be involved in providing tolerance to toxins produced by other phyllosphere colonizers, as seen recently with the clone expressing a putative glyoxylase/ bleomycin resistance dioxygenase showing neutralizing activity against toxoflavin, produced by *Burkholderia gladioli* (Choi et al., 2018). SGNH hydrolase-like domain containing acetyltransferase protein, AlgX, is involved in alginate biosynthesis in *Pseudomonas aeruginosa* and in virulence during cystic fibrosis. Although alginate is known to be involved in epiphytic fitness in *Pseudomonas syringae* (Yu et al., 1999), its contribution towards stress tolerance in xanthomonads remains to be investigated. Another gene belonging to PD-(D/E)XK nuclease superfamily, also characterized in contact-dependent DNA-degrading effectors (Sirias et al., 2020), was identified in both commensals and weak pathogens. Such nuclease effectors may impart a competitive advantage to the commensals and weak pathogens in a mixed species community as they establish their niche. At least three TonB-dependent receptors were identified to be associated with commensals and weak pathogens. Overall, TonB-dependent receptors are over-represented in xanthomonads and are thought to impart the adaptive ability to xanthomonads to utilize a wide variety of carbohydrates (Blanvillain et al., 2007). Another gene associated with commensal and weak pathogens belonged to the NRAMP (natural resistance-associated macrophage proteins) family, conserved from bacteria to eukaryotes. These proteins function as proton-dependent manganese transporters in bacteria and have evolved to adapt to oxidative environments (Cellier et al., 2001; Kehres et al., 2000). Enrichment of these proteins in commensals and weak pathogens may explain why they can be simultaneously isolated along with pathogens from infected tissue and why they can withstand host defenses, including reactive oxygen species.

We also screened the genomes for *fliC*, flagellin encoding gene, and specifically, the canonical flg22 epitope sequence to see the extent of variation within xanthomonads. It is not known whether commensal and weakly pathogenic xanthomonads can evade MAMP-triggered immunity induced by flagellin. Polymorphism in the 43^rd^ amino acid residue in the *fliC* sequence, a part of the flg22 epitope, was previously observed in pathogenic xanthomonads (Malvino et al., 2022). More specifically, a change from the canonical residue Val^43^ to Asp^43^ allowed pathogenic xanthomonads to escape detection by FLS2 in *Arabidopsis* and tomato (Malvino et al., 2022; Sun et al., 2006). We observed that all commensals and weak pathogens carried the canonical Val^43^ in the flg22 epitope, except for *X. maliensis* LMG27592. In addition, some pathogenic xanthomonads also had the immunogenic version of flagellin (Table S3). Some commensals and weak pathogens possessed multiple copies (up to 3) of *fliC* in their genome. Based on their similarity to the canonical flg22 epitope residues, they can be expected to possess immunogenic properties. However, variations in other parts of the gene may have a role in modulating immunity in various hosts.

We also identified genes that were enriched in pathogens compared to commensals. As expected, the majority of such genes were genes involved in T3SSs and effectors (Table S13, Figure S11). Other features enriched in pathogens included glycosyl hydrolases, phage proteins, and chemotaxis proteins. Certain phage-related proteins and transposases were also identified as exclusively present in pathogens but absent in commensals, indicating their role in mobilizing pathogen fitness and virulence genes. This finding suggests that it might be possible to devise phage-based control strategies using phages selective towards pathogens (Table S13). Pathogens also exclusively contained certain transcriptional regulators, genes involved in c-di-GMP signaling pathway and chemotaxis, lipid metabolism, and carbohydrate and amino acid metabolism compared to commensals (Table S14).

Genes that were enriched in pathogens compared to nonpathogens (commensals and weak pathogens) included a TonB-dependent receptor involved in Fe transport, a chemotaxis protein, specific cell-wall degrading enzymes, and an anti-sigma factor and AraC transcriptional regulator (Figure 5, Table S15A, S15B, and S15C). Among the genes depleted in commensals and weak pathogens was an adenylosuccinate synthetase (S-AMPS) involved in *de novo* purine biosynthesis in the cytoplasm. This pathway is connected to the TCA cycle and, thus, to the central metabolism. This enzyme links GTP hydrolysis to inosine monophosphate (IMP) condensation with L-aspartate to produce adenylosuccinate (S-AMP). It can be hypothesized that pathogens may experience limitations of these compounds in their niche, for example, apoplast or xylem, and, thus, have evolved the ability to synthesize S-AMP. This enzyme was found in the periplasm-enriched fraction of *X. citri* but not in *X. fuscans* subsp. *aurantifolii* type B (Zandonadi et al., 2020). Another gene involved in lipid metabolism, annotated as phosphatidylserine/phosphatidylglycerophosphate/cardiolipin synthase or related enzyme was associated exclusively with pathogenic *Xanthomonas*. *X. campestris* was characterized for its unique lipid metabolism pathways with a bifunctional enzyme for PE synthesis that functions in a serine-dependentor an ethanolamine-dependent pathway, depending upon the availability of substrates *in-planta* (Aktas & Narberhaus, 2015).

## Discussion

Since nonpathogenic *Xanthomonas* strains were first co-isolated with pathogenic strains or in close association with plants or plants debris, the interest in exploring the role of these strains in the evolution of the genus, their pathogenic potential, and their contribution towards the microbiota-mediated extended plant immunity have increased (Bansal et al., 2020, 2021; Cesbron et al., 2015; Entila et al., 2023; Essakhi et al., 2015; Gonzalez et al., 2002; T. Li et al., 2020; Martins et al., 2020; Pfeilmeier et al., 2021, 2023; Triplett et al., 2015; Vauterin et al., 1996). In this study, we attempted to address these aspects to understand how the diversity spanning the commensal to pathogenic lifestyles across the genus *Xanthomonas* has evolved. We harnessed the unexplored diversity that the collection sequenced in this study brought, particularly the previously poorly explored group 1 xanthomonads species (Studholme et al., 2011). These new genomes were combined with a collection of 1,834 high-quality publicly available *Xanthomonas* genomes to obtain a phylogenetically representative set of 337 genomes spanning the different lifestyles present in the genus for comparative analysis.

We observed variable, but not random, presence/absence of T3SS clusters as indicated in the previous studies (Cesbron et al., 2015; Fang et al., 2015; Merda et al., 2017; Triplett et al., 2015).

This study extends these findings to include some atypical T3SS clusters not limited to group 1 xanthomonads. This finding of gain of atypical T3SSs in addition to the *hrp2* cluster in some group 2 strains raises important questions about its functional significance and ecological role. The variable presence of T3SSs and effector repertoire sizes in xanthomonads ranging from 0 to 41 effectors indicate a plethora of diversity in the lifestyles of xanthomonads associated with plants along a continuum from being commensal endophytes to opportunistic pathogens or weak pathogens to full blown pathogens.

As a genus for which research has been highly focused on pathogenic potential, commensal or nonpathogenic xanthomonads represent a largely ignored component (Vauterin et al., 1996). Unlike pseudomonads, this side of the continuum that spans from commensal to opportunistic lifestyles has not been well studied in xanthomonads. We lack understanding as to what extent nonpathogenic xanthomonads play a role in being evolutionary partners of plants with adaptive value contributing to overall plant health or whether they are just the members that contribute to niche filling, a process influenced by plant traits but are of minor adaptive importance in terms of fitness or growth benefit to the plant host. Hacquard et al. (2017) proposed a system involving multiple layers of barriers for establishing homeostasis with plants. Here, we assess how xanthomonads may have established association with the plants across the continuum of lifestyles from commensals to pathogens using this framework. The first two protective layers are: (1) microbiota exhibiting nutritional and niche competition and (2) plant physical barriers. Based on the association analysis conducted in this study, we identified several traits in commensals, such as enrichment in type VI secretion system genes, transcriptional regulators, and carbohydrate metabolism genes, which may indicate their ability to overcome nutritional and niche competition with other microflora members associated with a wide range of plant hosts.

Distinct cell-wall degrading enzyme repertoire in commensals, weak pathogens, and pathogens may also indicate their differential ability to overcome epidermal cell barriers. The next layer in maintaining homeostasis with plants is MAMP-triggered immunity (MTI). We found that commensal, weakly pathogenic, and few pathogenic xanthomonads possess canonical flg22 immunogenic epitopes, indicating that plants recognize and mount an innate immune response against them. Whether they can suppress this defense response may depend on the presence of T3SSs and effectors. It is hypothesized that a minimal repertoire of effectors present in some nonpathogenic *X. arboricola* may help them suppress the MTI (Merda et al., 2017). Some commensal strains from our collection had no known effectors, indicating that MTI may explain their low abundance and lower in planta population. Their simultaneous isolation with pathogenic xanthomonads may also suggest that association with pathogens allows them to take advantage of innate immune response suppression performed by the pathogens. However, the ability of commensal and weakly pathogenic xanthomonads to activate MTI may also suggest their contribution towards extending plant immunity against other pathogenic bacteria or fungi, as seen in *Sphingomonas* (Innerebner et al., 2011). Apart from this epitope conservation, we observed variation in the rest of the flagellin sequence of commensals and weak pathogens. The importance of such sequence diversification in commensals and multiple copies of flagellin in evading MTI needs to be further explored. MTI suppression by type III secreted effectors has been demonstrated in many pathogenic xanthomonads. Analysis in this study showed that weak pathogens could also cross the MTI barrier due to the presence of a larger set of effectors (7-12 effectors) and intact T3SS. Whether a minimized ancestral T3E repertoire of commensal xanthomonads is sufficient for them to overcome the MTI barrier needs to be explored further as commensals lack T3SS and thus secretion and translocation of these effectors might be of question. However, Merda et al. (2017) indicated the possibility of the secretion of effectors, specifically *xopR* and *avrBs2,* mediated by the flagellar apparatus (Journet et al., 2005). These effectors may also have functions independent of T3SS. Simultaneous colonization of commensals and pathogenic xanthomonads may also mean that commensal xanthomonads coordinate the action of these effectors and share these effectors as a public good with the pathogenic members, similar to the phenomenon demonstrated with studies using effectorless *Pseudomonas* strains (Ruiz-Bedoya et al., 2023). Experiments evaluating the effect of coinfection on their collective virulence may further our understanding of the importance of reduced effector repertoires in commensal xanthomonads. These effectors in commensal strains may also have a role outside the host, similar to that shown for AvrBs1, IS476, and the associated plasmid offering enhanced overwintering potential (O’Garro et al., 1997). Further functional assessment of MTI-inducing and MTI-suppressing abilities of the commensals and weak pathogens from diverse clades across the phylogeny may clarify whether they can actively or passively cross the MTI barrier and how they may contribute towards microbiota-host homeostasis and ultimately towards plant growth-defense tradeoff and plant fitness (Ma et al., 2021). Although the opportunistic nature of nonpathogens has been documented (Gitaitis et al., 1987), the contribution of T3SS and an associated minimized effector repertoire or distinct CWDE repertoire towards such conditional pathogenicity has not been experimentally validated. Alternatively, conditional pathogenicity may result from altered host-microbiota homeostasis and a compromised immune response rather than the involvement of T3SS or CWDEs alone. This important question of the contribution of nonpathogenic xanthomonads as a member of the microbiota has been investigated by two independent studies that emphasized the role of T2SS and cell wall degrading enzymes in mediating the shift in microbiota and conditional pathogenicity (Entila et al., 2023; Pfeilmeier et al., 2023). Further, Entila et al. (2023) showed that conditional pathogenicity of a nonpathogenic xanthomonad strain, lacking *hrp2* cluster, is kept in check by suppression of cell-wall degrading enzymes by plant NADPH oxidase respiratory burst oxidase homolog D (RBOHD) through a negative feedback loop between DAMP-triggered immunity-led reactive oxygen species production by Arabidopsis and T2SS/cell-wall degrading enzymes.

The diversity of commensals and weak pathogens across the genus phylogeny present opportunities to study evolution of xanthomonads specifically in the context of important virulence factors such as T3SS, effectors, and CWDEs that explain their plant-associated lifestyles. While CWDEs have been proposed to be acquired by the ancestor of xanthomonads, they seemed to have diversified along the course of evolution as xanthomonads established associations with diverse hosts. The differential repertoire of CWDEs accompanied by unique TonB-dependent receptors among pathogens and commensals may explain the differential niche colonization. While we did not study patterns of gain/loss of specific CWDEs, we observed that commensal and weak pathogens have distinct sets of CWDEs compared to pathogens, similar to the observation with pathogenic and nonpathogenic *X. arboricola* (Cesbron et al., 2015). Different methyl-accepting chemotaxis proteins in commensals and pathogens may also suggest how the perception of the environment may be lifestyle-dependent. Similar to the observations by Merda et al. (2016, 2017), we found evidence for an ancestral gain of the *hrp2* cluster before the split of Group 2 xanthomonads. Group 1 xanthomonads have independently acquired different T3SS clusters. Thus, commensals belonging to Group 1 xanthomonads may possess ancestral traits that allowed them to closely associate with plants, such as diverse regulators, carbohydrate metabolism, and defense/repair-related genes. Subsequently, upon diversification of Group 2 xanthomonads into subsequent species, each displaying a high degree of host specificity, the effector repertoire reshuffling was observed (Hajri et al., 2009). Examining the genomic context of *hrp2* cluster and identities of core *hrp2* genes led us to hypothesize that replacement of *Xcc*-type *hrp2* cluster by *Xeu*-type *hrp2* cluster occurred within clade B strains, except in *X. nasturtii*, through rearrangements. Merda et al. (2016, 2017) also observed gene flow among T3SS and effectors of Group 2 xanthomonads. The subsequent loss of *hrp2* cluster in certain lineages representing commensal xanthomonads was observed (Figure 2). T3SS and associated effector loss may suggest that T3SS may impose a fitness cost. Thus, under this model, commensals belonging to Group 2 may have been derived by regressive evolution from pathogenic strains. We also assessed patterns of gain/loss of *hrp2* regulators, *hrpG/X*, analyzed in this study to identify clues to their involvement in the regulation of T3SS, associated limited repertoire in weak pathogens, or their existence in commensals in the absence of T3SS. Pfeilmeier et al. (2023) found that these master regulators do not regulate T2SS in nonpathogenic *Xanthomonas* strains lacking T3SS. This suggests a distinct regulatory network in play in commensal and pathogenic xanthomonads upon colonization of the host. Our analysis also indicated that many commensal and pathogenic strains engage in gene exchange and possess shared mobile genetic elements and carrier genes. Although experiments demonstrating the transfer of T3SSs from pathogenic to nonpathogenic xanthomonads were insufficient to impart pathogenicity phenotype to nonpathogenic strains (Meline et al., 2019), gene flow among certain strains may explain the re-acquisition of T3SS. Such gain/loss events of T3SS and associated effectors may explain the heterogenous distribution of the *hrp2* cluster and diversity in effector repertoires across strains. Merda et al. (2016) showed that genetic exchange involving life history traits between pathogenic and nonpathogenic *Xanthomonas* strains occurred likely through horizontal gene transfer and suggested a possibility of nonpathogenic strains acting as reservoirs of traits that allow the emergence of novel pathogenic strains (Meline et al., 2019). Here, we gathered empirical evidence for the transfer of an IS element from the nonpathogenic to the pathogenic strain upon co-inoculation of the same plant. This exchange disrupted the avirulence gene in the pathogen under selection pressure allowing it to overcome host resistance. This finding further highlights the role of commensals as a reservoir of traits that may contribute to the adaptation of pathogens resulting in new outbreaks, as demonstrated by Lee et al. (2020). Also, HGT between pathogen and commensal strains has been demonstrated in some bacteria, converting nonpathogenic strains into pathogenic ones (Brouwer et al., 2013). Such adaptive traits may have been subject to gene transfer among commensal and pathogenic strains.

However, this study also demonstrates that T3SS and effectors are not the sole lifestyle determinants in xanthomonads. Commensals and weak pathogens have evolved strategies for tolerance to stresses with distinct sets of chemotaxis proteins, type VI secretion systems, TonB-dependent receptors, chaperones, and transcriptional regulators. We previously examined gain/loss patterns of type VI secretion system clusters across the *Xanthomonas* phylogeny and found evidence for non-random acquisition of T6SS and gene flow in core genes and effectors (Liyanapathiranage et al., 2022). Further examination of gain/loss patterns of traits enriched in non-pathogens may help assess support for the regressive evolution model to explain the origin of commensal strains.

As for the strains isolated from rainfall, some *X. arboricola* strains cluster phylogenetically with pathogenic strains of the same species isolated from plants and are indistinguishable in regard to the presence of a T3SS and the size of their effector repertoires. This suggests that they are pathogens that were simply caught while migrating through the atmosphere between host plants.

The rainfall isolates identified as *X. euroxanthea* and as members of the novel species cluster VIII, are very similar to strains without T3SS and a very small number of effectors previously isolated from plants. They can thus also be expected to generally live in association with plants, although as non-pathogens, and they were also isolated from the atmosphere while being dispersed from one plant to the other. The situation may be similar for the four non-pathogenic strains isolated from rainfall that belong to the novel species clusters V and VII. However, since no closely related strains from plants are yet known, a life style independent of plants cannot be excluded.

Finally, systematic evaluation of genomic traits associated with lifestyles might guide us as we develop diagnostic strategies to differentiate pathogenic from nonpathogenic strains associated with seed samples or infected field samples. A recent machine-learning approach developed to predict the phenotype of plant-associated xanthomonads indicated specific domains associated with pathogenic and nonpathogenic lifestyles (Molder et al., 2021), many of which were found to overlap with the candidates identified in this study. Our study further confirms that diagnostic methods cannot rely on type III secretion system gene markers alone to identify pathogenic xanthomonads.

This study highlights the diversity of lifestyles across the genus phylogeny along the continuum of commensal, weakly pathogenic, and pathogenic strains. We find that type III secretion system and effectors are not the only factors that define these lifestyles. Several niche adaptative factors were identified to be associated with each lifestyle. Commensals establish themselves on different hosts in the presence of various host defenses and competing microflora, as well as derive complex nutrients from a wide range of hosts while sustaining their populations on a broad range of hosts. We also observed distinct cell-wall degrading enzyme repertoires that distinguish pathogenic vs. commensal or weakly pathogenic lifestyles. Conversely, pathogens rely on type III secretion system and associated effectors to subvert defense responses. Overall, commensals possess genes that allow them to tolerate stresses rather than avoid them in the absence of T3SS.

## Author Contributions

The study was conceptualized and designed by NP, MAJ, BAV, and JBJ. Strains were collected from various sources by NP, BAV, JBJ, and MAJ. RB and EN processed the samples and conducted greenhouse experiments. TW assisted with sequencing and data availability from JGI. The data was analyzed by RB, MMP, RMB, KW, NW, TP, and NP. Specifically, MMP conducted OrthoFinder analysis, taxonomic placement, pathogenicity assay, T3SS, and T3E analysis, and wrote the initial draft of the manuscript. NW and TP conducted GLOOME analysis. RB analyzed CWDEs. KW performed MGEFinder analysis. NP carried out association analysis. RMB assisted with constructing heatmaps for association analysis. RB formatted figures and tables. The final manuscript was written by RB and NP, with input from all the other co-authors.

## Acknowledgments

The work (proposal: 10.46936/10.25585/60001156) conducted by the US Department of Energy Joint Genome Institute (https://ror.org/04xm1d337), a DOE Office of Science User Facility, is supported by the Office of Science of the US Department of Energy operated under Contract No. DE-AC02-05CH11231.

We thank CIRM-CFBP (Beaucouzé, INRAE, France http://www6.inra.fr/cirm_eng/CFBP -Plant-Associated-Bacteria) for strain preservation and supply. We also thank Abi S. A. Marques and Marisa A. S. V. Ferreira for participating in the strain sampling from bean.

## Data availability

Sequence data generated from this work have been deposited in the DOE JGI and GenBank database under the JGI taxon id and Bioproject accession number as given in Table S1.

