## Supplementary figures for "Genetic and functional diversity help explain pathogenic, weakly pathogenic, and commensal lifestyles in the genus *Xanthomonas*"

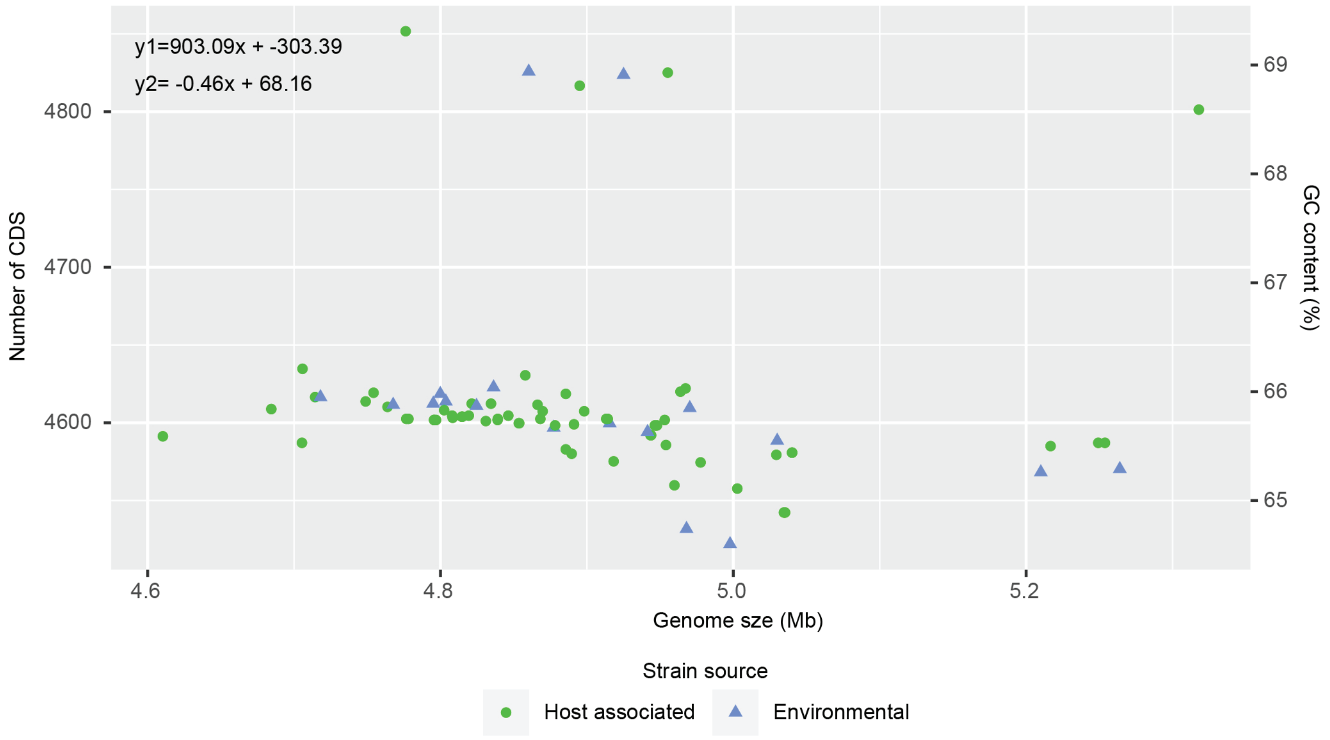


**Figure S1:** Relationship between genome size, GC content, and number of CDS for the genomes sequenced for this study. *Xanthomonas* sp. 60 is excluded from the plot.

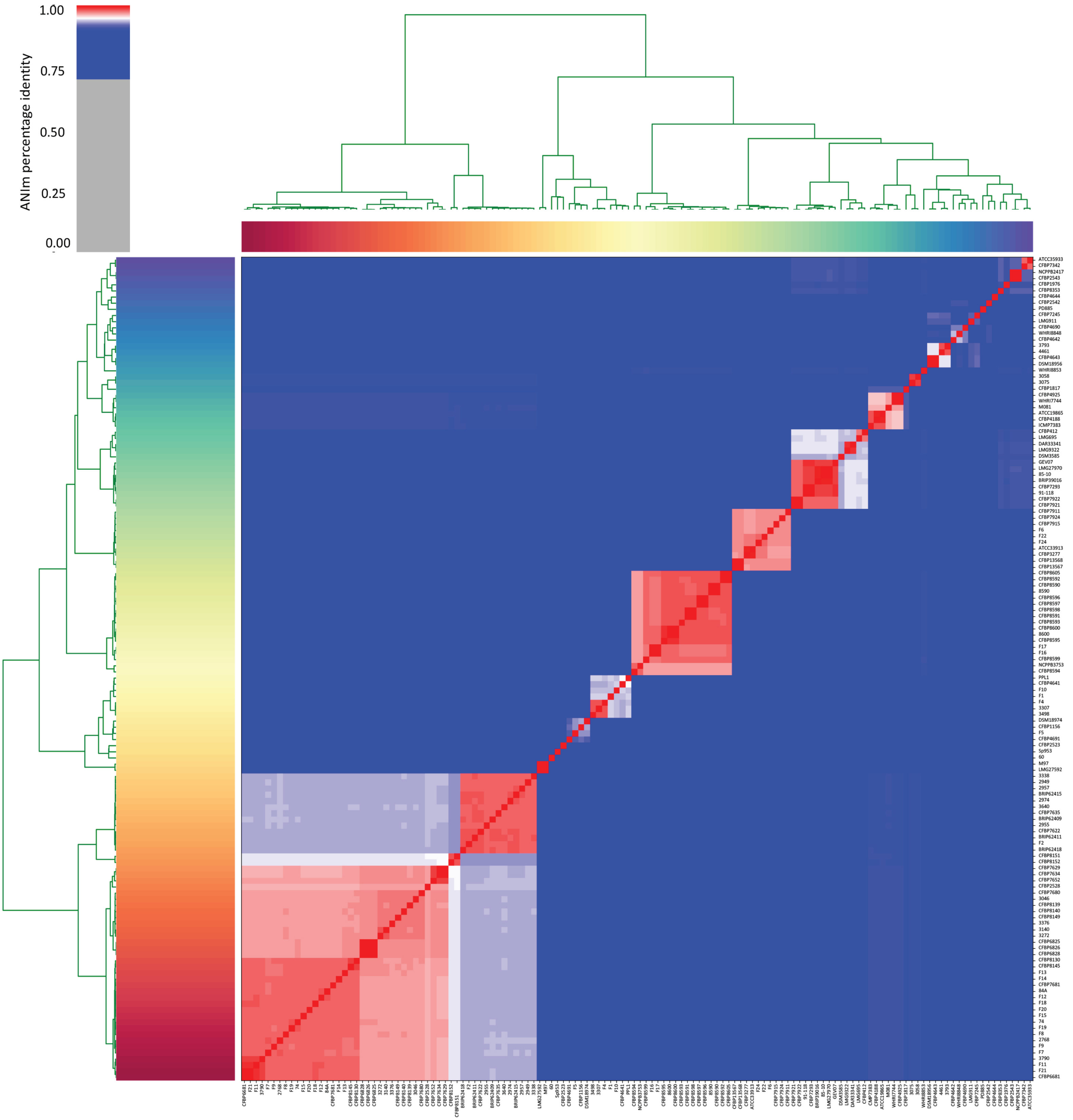


**Figure S2:** Average nucleotide identity (ANI)-based heatmap showing the status of the 134 *Xanthomonas* strains. The intensity of the color indicates the level of identity of all-versus-all genomes as depicted by the scale.

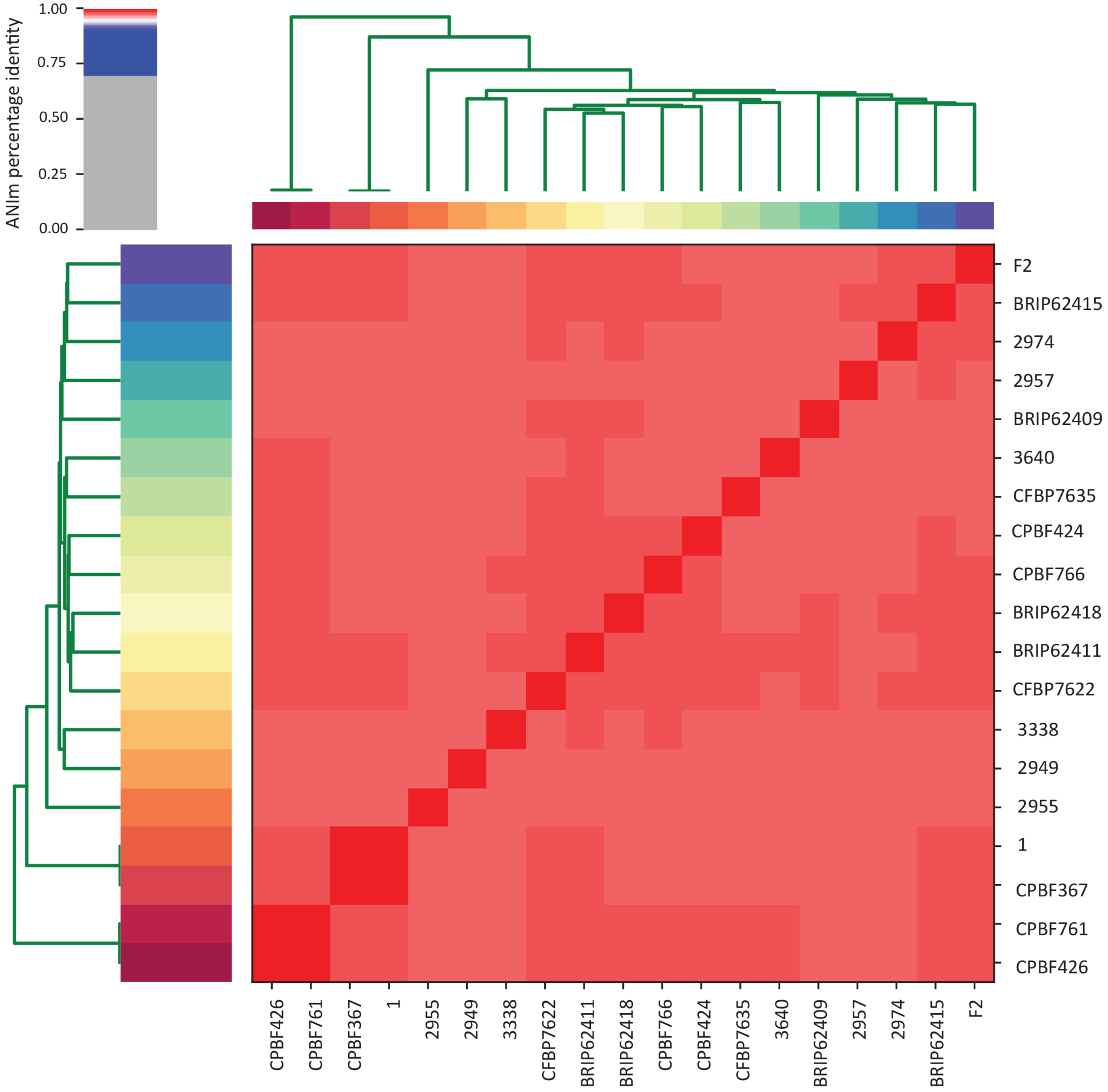


**Figure S3:** Average nucleotide identity (ANI)-based heatmap showing the status of the representative *Xanthomonas euroxanthea* strains and strains from this study. The intensity of the color indicates the level of identity of all-versus-all genomes as depicted by the scale.

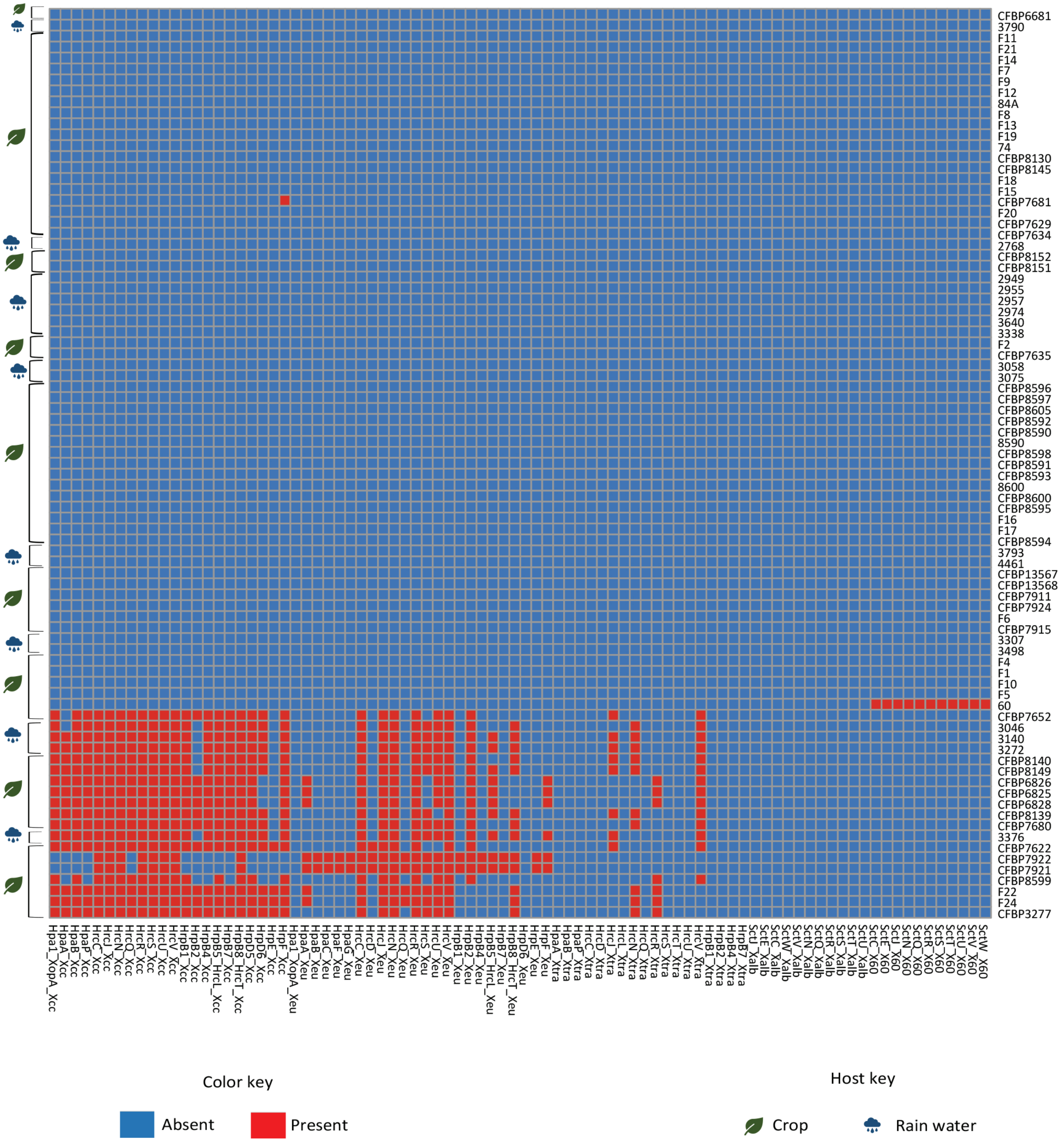


**Figure S4**: Heatmap showing the status of T3SS genes in *Xanthomonas* strains used in this study. Here, query genes names from the different species are indicated as gene_*Xeu* (genes from *Xanthomonas campestris* pv. *vesicatoria* 85-10), gene_*Xcc* (genes from *Xanthomonas campestris* pv. *campestris* ATCC33913), gene_*Xtra* (genes from *Xanthomonas translucens* pv. *translucens* DSM18974), gene_*Xalb* (genes from *Xanthomonas albilineans* CFBP 2523), and gene_*X60* (genes from *Xanthomonas* sp. 60). Red color represents the presence while blue color represents the absence of the gene.

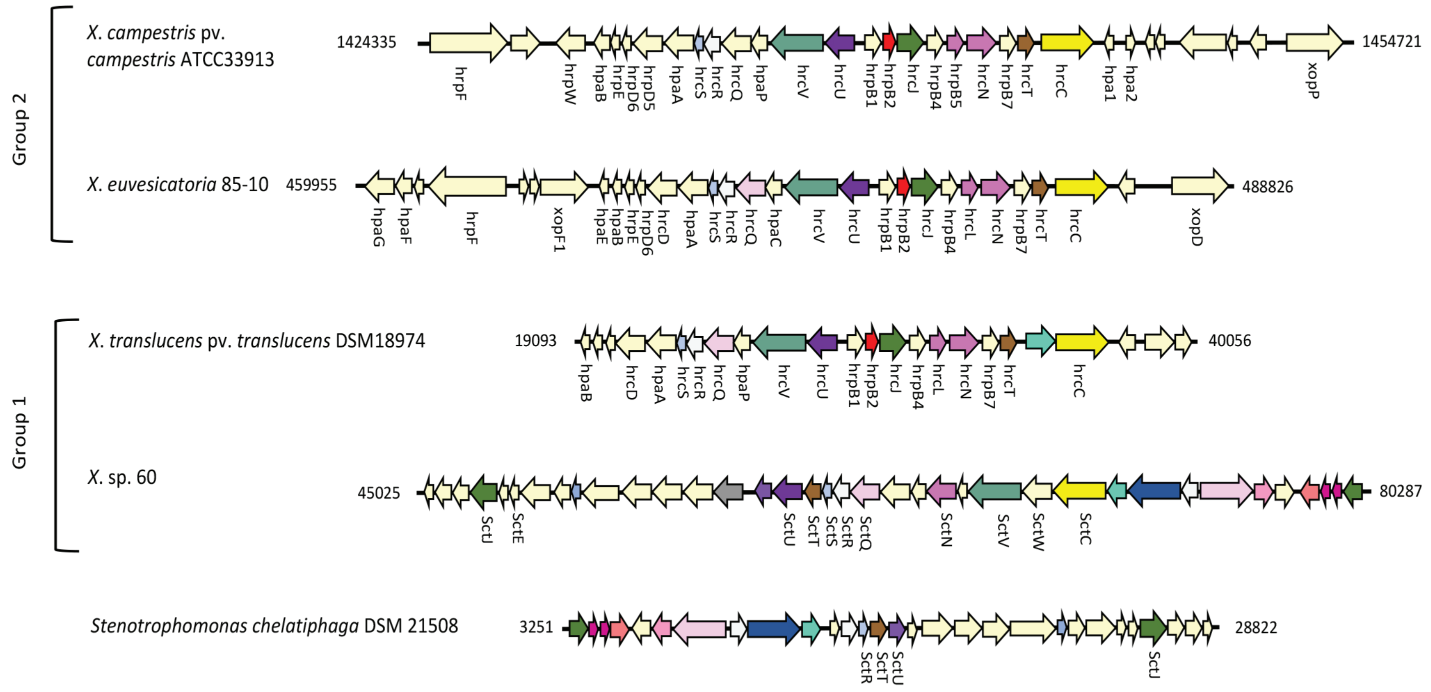


**Figure S5**: Schematic representation of the genetic organization of T3SS clusters found in *Xanthomonas campestris* pv. *campestris* ATCC33913 (*Xcc*), *Xanthomonas campestris* pv. *vesicatoria* 85-10 (*Xeu*), *Xanthomonas translucens* pv. *translucens* DSM18974 (*Xtr*), *Xanthomonas* sp. 60, and *Stenotrophomonas chelatiphaga* DSM 21508. Genes of the same color (except light yellow, which has no COG assignment) are from the same orthologous group. The gene names are according to the Sct T3S injectisome nomenclature and annotation available in the IMG/M system (https://img.jgi.doe.gov/cgi-bin/m/main.cgi).

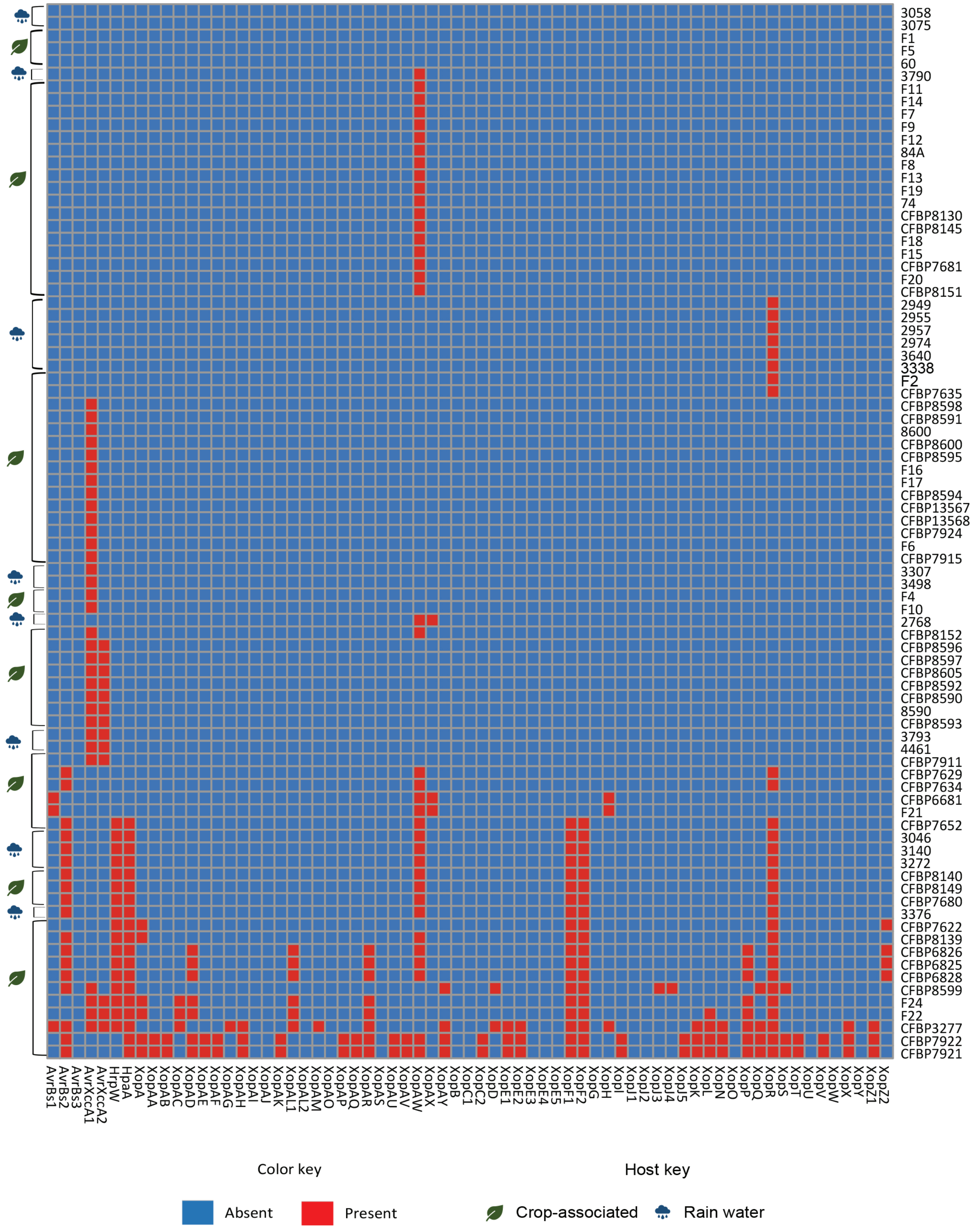


**Figure S6:** Heatmap showing the status of T3E in *Xanthomonas* strains used in this study. Here the query represents all effectors families, putative effectors (AvrBs1, AvrBs2, AvrBs3, Xop, HpaA, HrpW, and AvrXccA), and their diversity. Red represents the presence, while blue represents the absence of the gene.

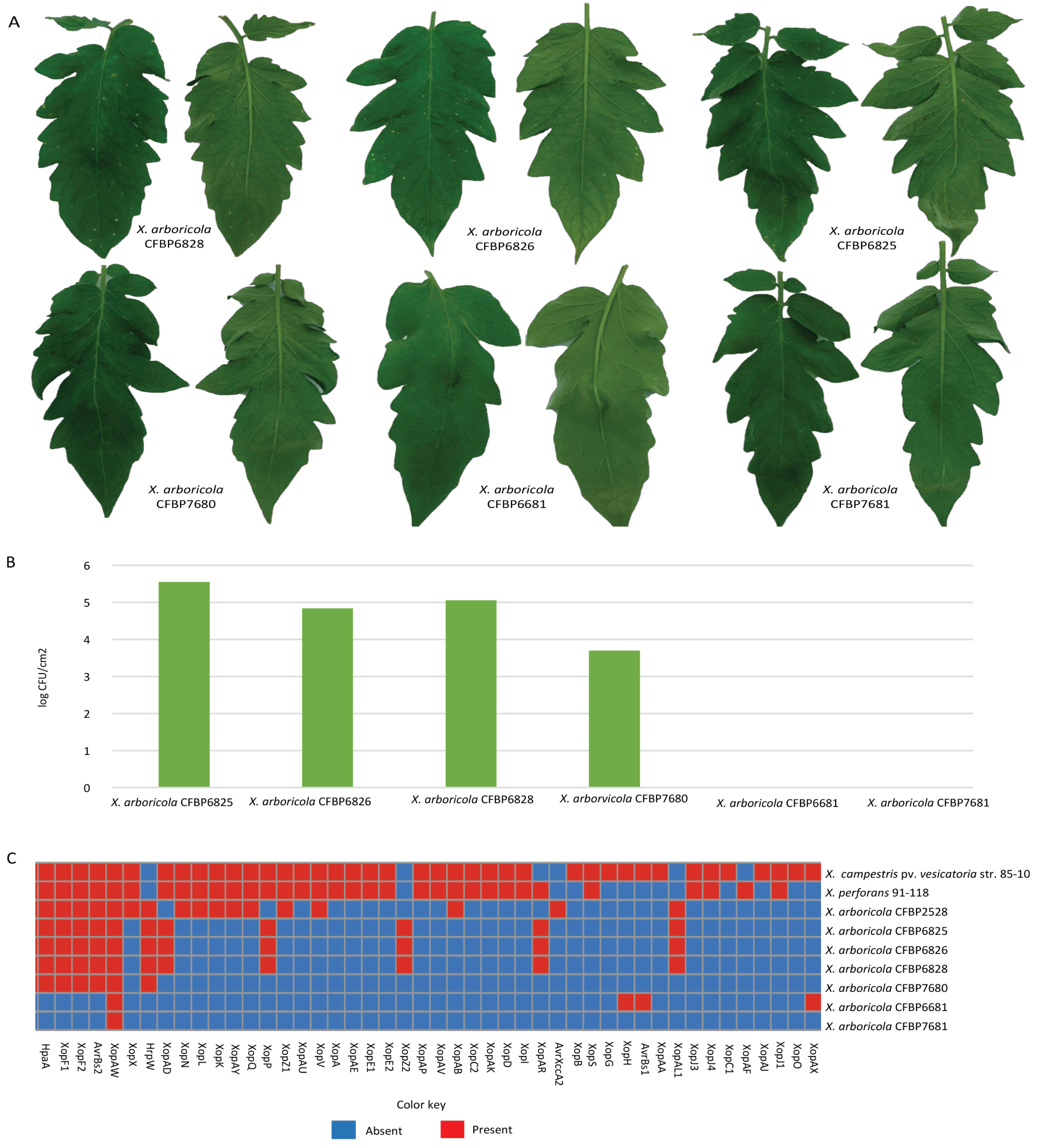


**Figure S7:** (A) Pathogenicity and (B) *in-planta* population and (C) distribution of T3E in *X. arboricola* strains (CFBP 6825, CFBP 6826, CFBP 6828, CFBP 6681, CFBP 7681, and CFBP 7680) on 4-5-week-old tomato cv. FL 47R 10 DAI (days after inoculation). The presence of common T3Es across different *Xanthomonas* strains is shown in red, while blue represents the absence of T3Es.

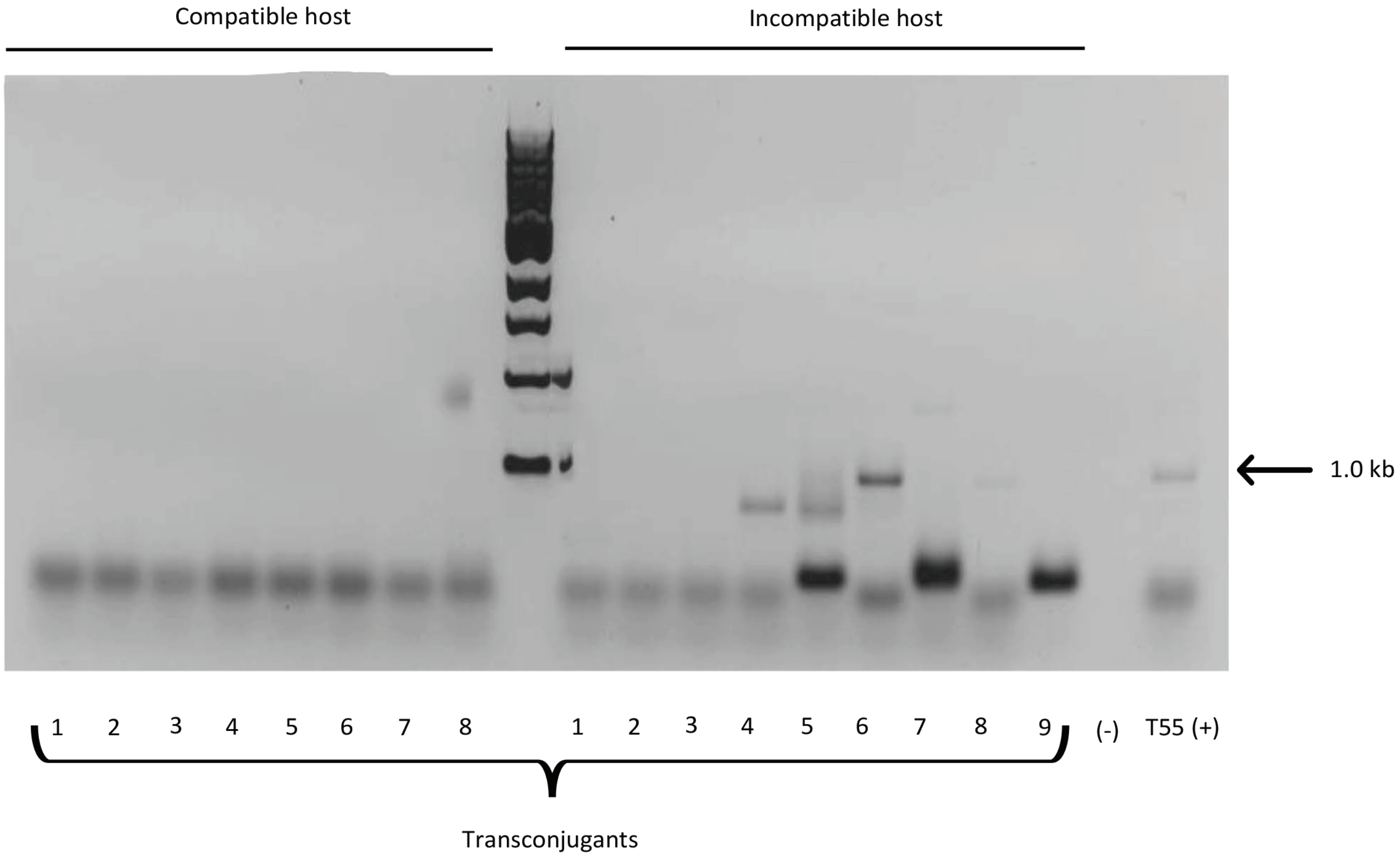


**Figure S8:** Analysis of the presence of type three secreted effector *avrBs1* with a transposon insertion among 50 randomly selected transconjugants. Mating experiments were conducted in both compatible (*Capsicum annuum* cv. Early California Wonder) and incompatible (*Capsicum annuum* cv. Early California Wonder 10R) backgrounds with *Xanthomonas euvesicatoria* strain 85-10 and *Xanthomonas* sp. strain T55. Each lane represents the pooled DNA of five to six individual transconjugants. The positive control (*Xanthomonas* sp. T55) shows an approximately 1.0 kb band indicative of *avrBS1* with a transposon insertion, transconjugants isolated from the incompatible host in lanes six and eight have acquired the disrupted avirulence gene.

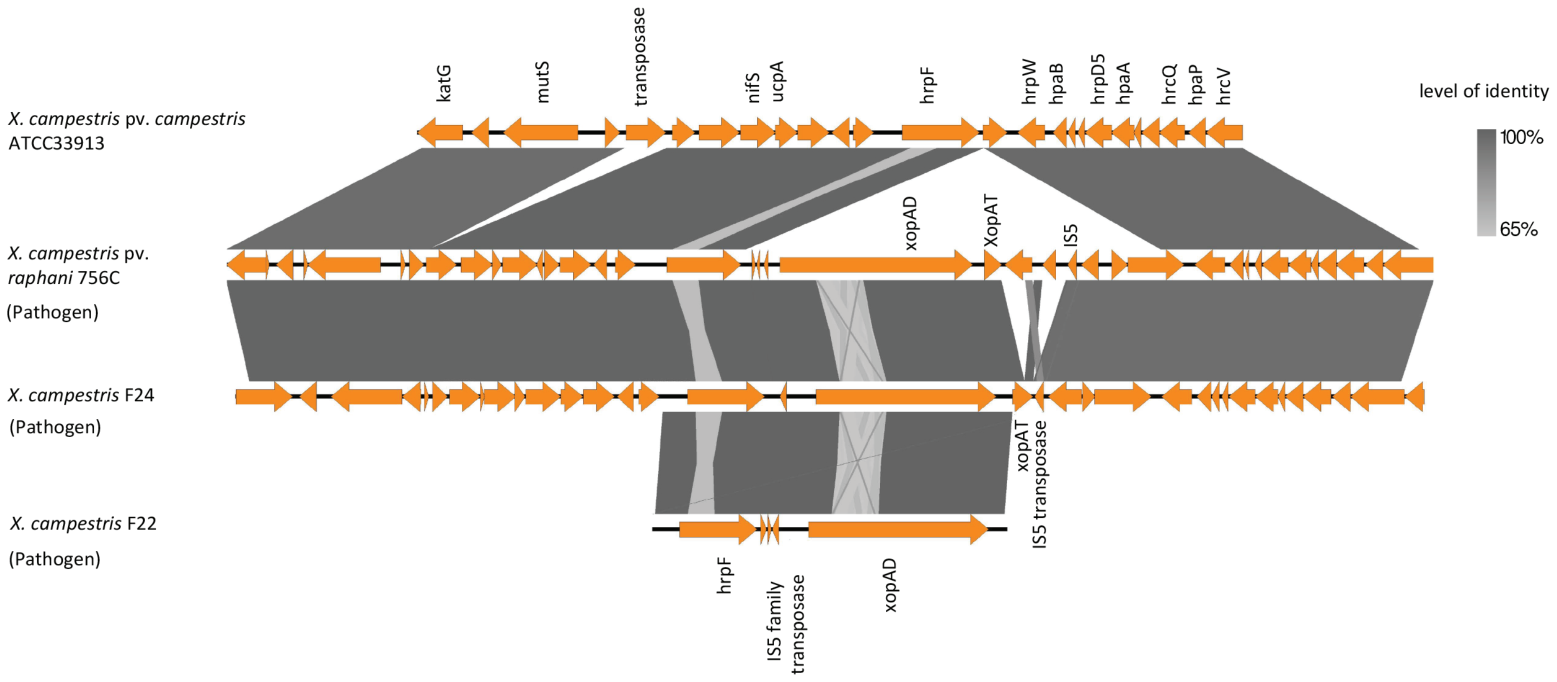


**Figure S9**: Schematic representation and comparison of the genetic organization of the mobile genetic element island identified by MGEfinder in *Xanthomonas campestris* pv. *raphanin* 756C, *Xanthomonas campestris* F24, and *Xanthomonas campestris* F22, highlighting the XopAD effector flanked on both ends by IS5 transposases.

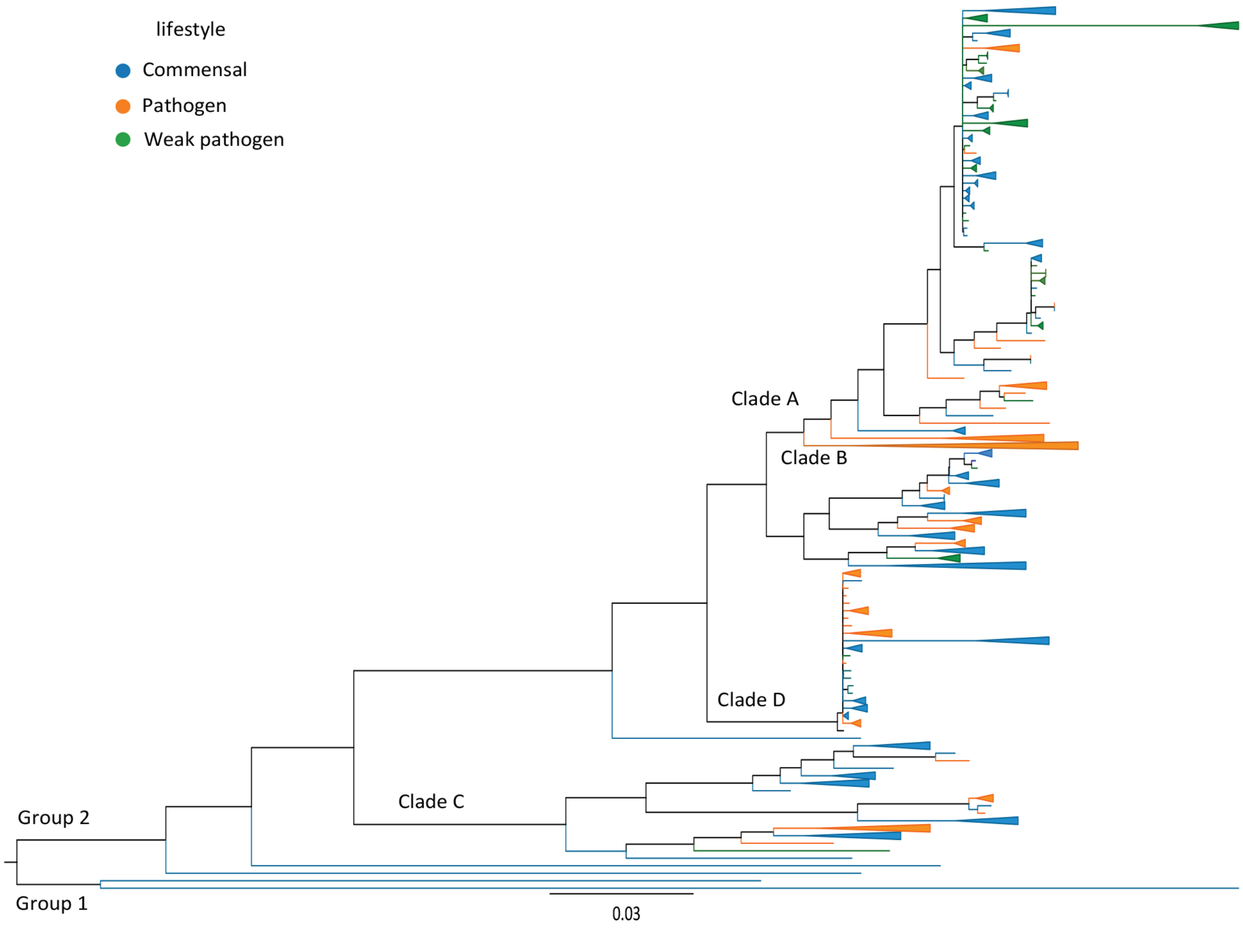


**Figure S10**: Maximum-likelihood phylogeny based on the orthogroups of 337 *Xanthomonas* strains. Commensal *Xanthomonas* strains are highlighted in blue branches, weakly pathogenic strains in green branches, and pathogenic strains in orange. Branches were collapsed to visualize better strain diversity and phylogenetic placement with different lifestyles.

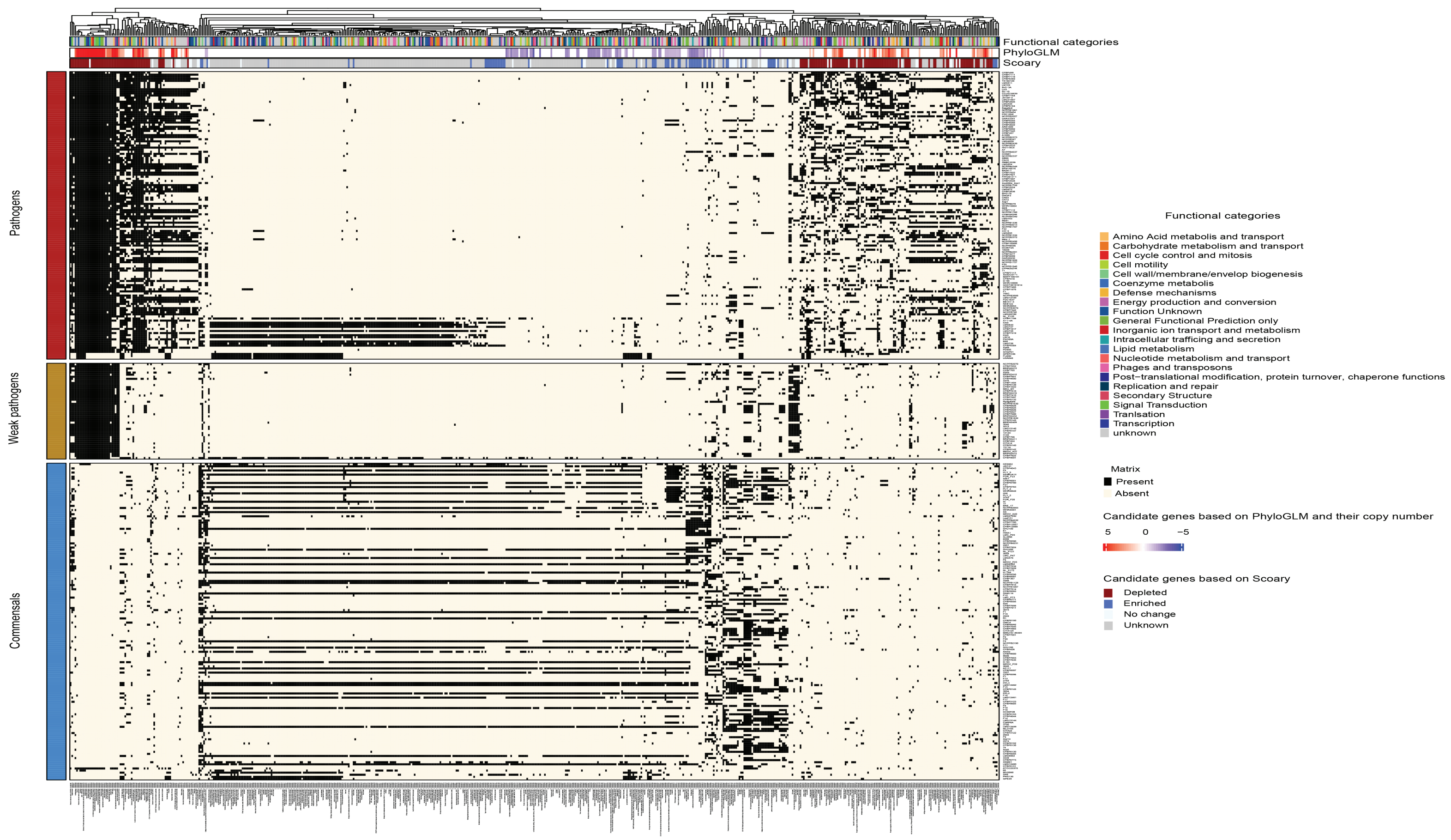


**Figure S11:** A complex heatmap showing results from association analysis correlating pathogenic, weakly pathogenic, and commensal phenotypes to the orthologs identified using Orthofinder. The functional categories are indicated for each candidate gene identified based on the intersection of three methods PhyloGLM, Scoary, and hyperglm (gene presence/absence), and those identified by PhyloGLM based on gene copy number.
